# CAPRIN1 localizes to CD63-positive MVB-like SARS-CoV-2 egress compartments and limits cytopathic effects

**DOI:** 10.64898/2026.08.25.746949

**Authors:** Joanna S. Woloszyn, Jonas Schroeder, Julia M. Berger, Elena Kotova, Robin Schäper, Susanne Pfefferle, Charlotte Uetrecht, Timothy K. Soh, Jens B. Bosse

**Affiliations:** Hannover Medical School, Institute of Virology, Hannover, Germany; Centre for Structural Systems Biology, Hamburg, Germany; Cluster of Excellence RESIST (EXC 2155), Hannover Medical School, Hannover, Germany; German Center for Infection Research (DZIF), Partner Site Hannover-Braunschweig, Hannover, Germany; CSSB Centre for Structural Systems Biology, Deutsches Elektronen-Synchrotron DESY & Leibniz Institute of Virology (LIV) & University of Luebeck, Notkestraße 85, 22607 Hamburg, Germany; University of Luebeck, Institute of Chemistry and Metabolomics, Ratzeburger Allee 160, 23562 Luebeck, Germany; Institute of Medical Microbiology, Virology and Hygiene, University Medical Center Hamburg-Eppendorf (UKE), Hamburg, Germany; Department of Virology, Bernhard-Nocht Institute for Tropical Medicine, Hamburg, Germany

**Keywords:** SARS-CoV-2, expansion microscopy, CAPRIN1, G3BP1, MVB, CD63, viral egress, cell death, syncytia

## Abstract

Stress granules (SGs) are cytoplasmic condensates that assemble under cellular stress, including viral infection. The best-characterized evasion strategy in SARS-CoV-2 infection involves the viral nucleocapsid (N) protein hijacking the SG core protein G3BP1, thereby preventing SG assembly. By performing network analysis of N protein interactomes, we identified CAPRIN1 as an underexplored candidate, despite its central role alongside G3BP1 in SG assembly. Here, we provide the first high-resolution spatial and functional data on CAPRIN1 in SARS-CoV-2 infection. Using ultrastructure expansion microscopy (U-ExM), we localized CAPRIN1 and viral egress markers to enlarged, CD63- and LAMP1-positive, multivesicular body (MVB)-like egress compartments, refining the current model of SARS-CoV-2 egress. CAPRIN1 was found on both the limiting membrane of these compartments and within the intraluminal vesicles, a pattern distinct from G3BP1. Knockout of CAPRIN1 increased infection-associated cell death, promoted an aberrant cell-death phenotype, and enhanced syncytia formation. Together, our data reveal distinct activities of CAPRIN1 during infection and identify the egress compartments as MVB-like, raising the possibility that SARS-CoV-2 repurposes more than one cellular pathway for egress.

## Introduction

Severe acute respiratory syndrome coronavirus 2 (SARS-CoV-2) is the causative agent of COVID-19. Among its four structural proteins, spike (S), nucleocapsid (N), membrane (M), and envelope (E) proteins, the N protein is the most abundant viral protein in an infected cell and the most functionally versatile, contributing to both viral processes and host cell modulation^1,2^. On the viral side, its central role is to bind the newly synthesized positive-sense viral RNA and package it into the ribonucleoprotein (RNP) complex at the double-membrane vesicles (DMVs), the replication organelles formed from remodelled rough endoplasmic reticulum (rER) membranes. The RNP complex then relocates to the ER-Golgi intermediate compartment (ERGIC), where N coordinates virion assembly through its interaction with the M protein^3,4^. Once assembled at the ERGIC compartment, newly synthesized virions must traffic out of the cell. Rather than using the conventional secretory pathway through the trans-Golgi network, β-coronaviruses were proposed to egress via lysosomal exocytosis^5^.

This exocytosis route depends on LAMP1-positive compartments and is actively shaped by the viral protein ORF3a, which sequesters VPS39 to disrupt Homotypic fusion and Protein Sorting (HOPS) complex-dependent membrane fusion, thereby preventing the compartments from maturing into lysosomes^6,7^. However, these compartments have been mainly characterized by LAMP1, complemented by markers such as Rab7 and cathepsin D^5,7,8^, all of which are shared across the late-endosome to lysosome continuum^9,10^. Whether they correspond to lysosomes or to related compartments, such as multivesicular bodies (MVBs), has therefore not been firmly established.

On the cellular side, N actively modulates the host cell, suppressing innate-immune signalling and interfering with stress granule (SG) formation^2,11^. Among its many reported host interactors, G3BP1 stands out: across published immunoprecipitation-mass spectrometry (IP-MS) datasets, it is the most consistently identified^12–14^, and it is also the best understood, with the N-G3BP1 interaction being structurally defined by N binding the G3BP1 NTF2-like domain^15^. Through this interaction, N hijacks G3BP1 to prevent SG condensation, its best-characterized effect on the host anti-viral response.

G3BP1 is the central nucleator of SGs, membraneless cytoplasmic condensates of mRNA, RNA-binding proteins, and stalled translation-initiation complexes that assemble when the cell is exposed to stress, including viral infection^16^. SGs contribute to anti-viral defense by sequestering viral RNA and concentrating innate-immune sensors such as protein kinase R (PKR)^17^. SARS-CoV-2 counteracts this defense: N hijacks G3BP1 and recruits it to viral replication-transcription complexes^18^. This interaction has been linked to pro-viral activity: an inhibitor designed to disrupt the N-G3BP1/2 interface dampens SARS-CoV-2 infection in cell culture^19^ and G3BP1 conditional knockout mice develop higher lung viral loads than wild type animals^20^.

Given the clear interaction between N and the SG pathway, other N interactors beyond G3BP1 may influence infection. One such candidate is cytoplasmic activation/proliferation-associated protein 1 (CAPRIN1), which promotes SG condensation, contributes to PKR-dependent innate-immune signalling, is a core partner of G3BP1, and binds the same G3BP1 NTF2-like domain targeted by N^17,21,22^. Despite this central position within the SG network, CAPRIN1 has been neither spatially characterized during SARS-CoV-2 infection nor tested for its impact on infection outcome.

Here, we provide the first spatial characterization in high resolution of the CAPRIN1-N protein interaction and reveal the functional consequences of CAPRIN1 loss. By co-staining infected cells for CD63 in addition to LAMP1 and combining it with ultrastructure expansion microscopy (U-ExM), we showed that the compartments at which viral egress markers accumulate were CD63-positive and MVB-like, refining the current model of β-coronavirus egress. Within these compartments, CAPRIN1 localized to both the limiting membrane (*i.e.* the membrane separating the compartment from the cytoplasm) and the lumen of the compartment, a distribution distinct from G3BP1. Additionally, loss of CAPRIN1 increased the frequency of cell death, promoted a morphologically aberrant cell-death phenotype, and enhanced syncytia formation. Together, these data demonstrate that CAPRIN1 plays a role in SARS-CoV-2 infection that is distinct from G3BP1 and influences the outcome of the infection at a stage not accounted for by its canonical role in stress granule biology.

## Materials and Methods

### Cells

Vero E6 cells (ATCC, Cat. No. CRL-1587) and Lenti-X 293T cells (Takara Bio, Cat. No. 632180) were cultured in Dulbecco’s Modified Eagle Medium (DMEM) with GlutaMAX (Thermo Fisher Scientific, Cat. No. 31966021) supplemented with 10% FBS Good (PAN-Biotech, Cat. No. P40-37500) under standard conditions (37°C, 5% CO₂).

A549-ACE2-TMPRSS2 (A549) cells (a kind gift from Krzysztof Pyrć^23^, Toni Luise Meister, and Stephanie Pfänder^24^) were cultured in DMEM/F-12 with GlutaMAX (Thermo Fisher Scientific, Cat. No. 31331028) supplemented with 10% FBS Good under standard conditions (37°C, 5% CO₂) and in the presence of 0.5 µg/mL puromycin (Invivogen, Cat. No. ant-pr-1) and 10 µg/mL blasticidin (Invivogen, Cat. No. ant-bl-05). The CAPRIN1-knockout A549 (CAPRIN1 KO) cells were derived from this line and maintained under identical conditions.

### Viruses

For all experiments except those using the KO cell line, we used a recombinant SARS-CoV-2 (rWuhan)^25^ generated by BAC-based reverse genetics from the Wuhan-Hu-1 backbone (pCC1-4k-Wuhan-Hu-1 nanoBAC based on GenBank NC_045512.2), carrying NanoLuciferase in-frame after the C-terminus of the ORF7a protein using a foot-and-mouth disease virus (FMDV) 2A linker^26^. Infection experiments with the CAPRIN1 KO cells were performed with the SARS-CoV-2 field isolate (HH-1) SARS-CoV-2/human/DEU/HH-1/2020 (GenBank accession no. MT318827.1). The phenotypes in this manuscript were compared between rWuhan and HH-1 (Supplemental Figure 1).

**Supplemental Figure 1:**
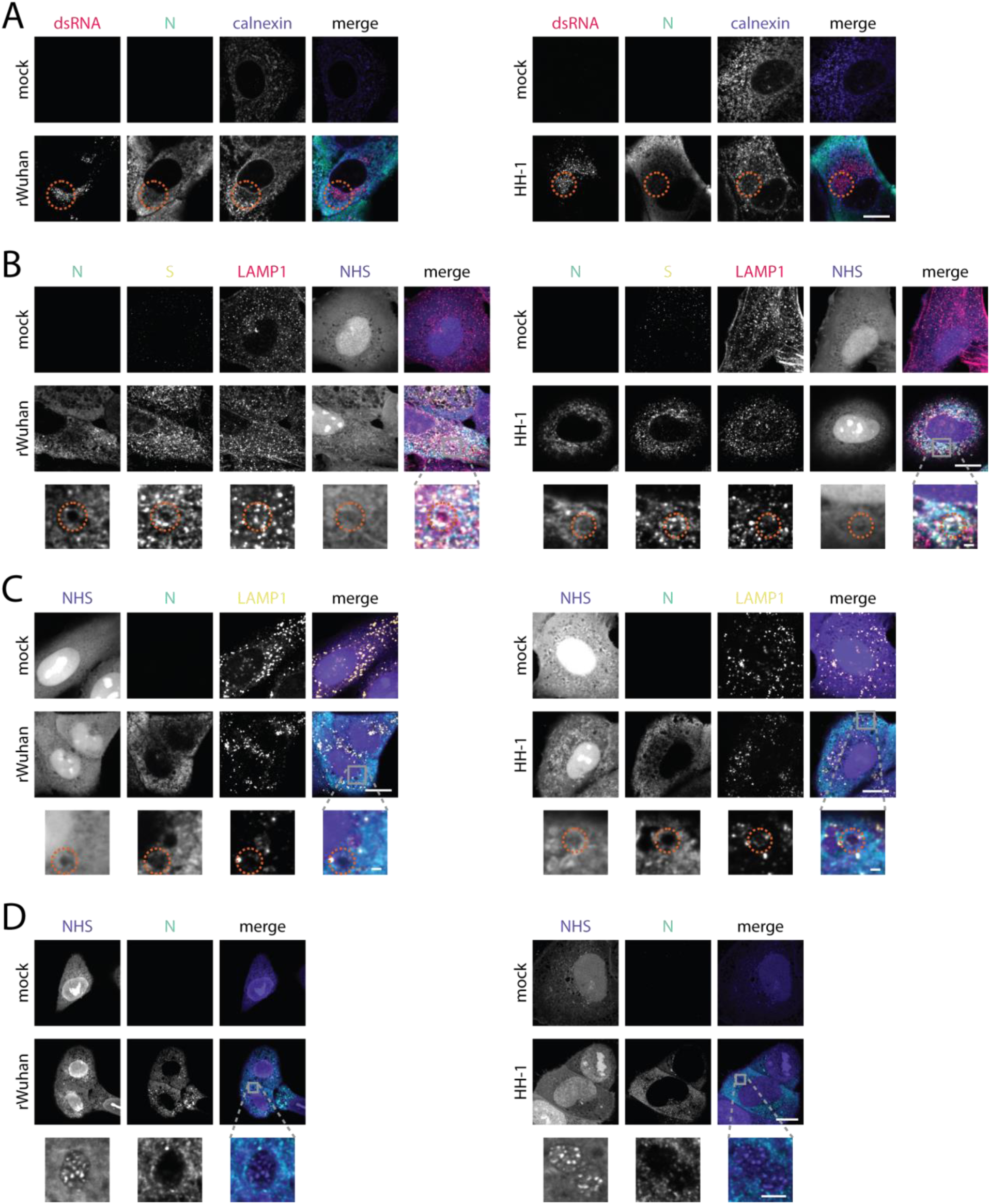
The recombinant SARS-CoV-2 strain Wuhan (rWuhan) and the field isolate strain HH-1 produce comparable replication- and egress-site phenotypes in infected cells. Cells were infected with the indicated virus at an MOI of 5 and fixed at 11 hpi. Scale bars, 10 µm; Zoom-in scale bars, 1 µm. For U-ExM images, the scale bars are post-correction. (A) Analogous to Figure 2B, infected cells were stained for dsRNA (red), N (green), and the ER marker calnexin (purple) to identify DMV replication sites (red circle). (B) Analogous to Figure 2E, infected cells were stained for N (green), S (yellow), NHS (purple), and LAMP1 (red) to identify egress compartments (red circle). (C) Analogous to Figure 4C, infected cells were stained with NHS (purple), N (green), and an alternate LAMP1 antibody (yellow) to identify egress compartments (red circle). (D) Analogous to Figure 4B, infected cells were processed for U-ExM and stained for NHS (purple) and N (green).

### Biosafety

All experiments involving infectious SARS-CoV-2 were performed under biosafety level 3 (BSL-3) conditions at the Bernhard Nocht Institute for Tropical Medicine (Hamburg, Germany), with genetic modification work and the handling of recombinant viruses approved by the responsible Hamburg authority (approval no. AZ I14-29/2022). Infected cells were fixed with 4.5% formaldehyde (SAV Liquid Production GmbH) in PBS for at least 30 min at room temperature (RT) to inactivate the virus. The inactivation protocol was validated and approved by the biosafety committee of the Bernhard Nocht Institute for Tropical Medicine.

### Network analysis

The mapping of the SARS-CoV-2 N interactome onto the human interactome (PICKLE human interactome, release 3.3^27^) was performed with a custom Python script using the python-louvain package. Python code is available at GitHub: https://github.com/QuantitativeVirology/SARS-CoV-2_N_CAPRIN1. Visualization of the networks was implemented with Gephi 0.10.1. The particularly dense regions of the human interactome were determined via the Louvain clustering method. Inside each cluster, the proportion of query proteins was determined and only those clusters with a query protein content above a threshold of 10% were extracted for a subgraph. The whole procedure was repeated on the subgraph with a threshold of 20% to yield a sub-subgraph, which could potentially contain the N-protein interactome. Louvain clustering is heuristic and can yield variable results depending on the initial conditions. To achieve a more representative result, the first and second clustering were performed 20 times each and a consensus clustering was aggregated.

### Immunoprecipitation (IP) of N

Each 15 cm dish of Lenti-X 293T cells was co-transfected with 20 µg STREP-TEV-N and 1 µg transfection control (pcDNA3 mCherry) expression plasmids using 120 µL PEI (Polysciences, Cat. No. 23966-2) in 3 mL Opti-MEM (Gibco, Cat. No. 31985070). Cells transfected with the corresponding empty vector served as a negative control. Two days post-transfection, cells were harvested, washed three times with PBS and lysed in 2 mL Lysis Buffer (Chromotek, Cat. No. lysbuf-30) for 40 min at 4°C under constant rotation.

Lysates were diluted with 2 mL Dilution Buffer (Chromotek, Cat. No. dilbuf-10) and incubated with 150 µL MagStrep Strep-Tactin XT beads (IBA Lifesciences, Cat. No. 2-5090-002) for 2 h. The beads were subsequently washed three times with wash buffer (IBA Lifesciences, Cat. No. 2-1003-100). Bound proteins were eluted by boiling in 1× SDS loading buffer (50 mM Tris pH 6.8 (Carl Roth, Cat. No. 4855.1), 2% SDS (Sigma-Aldrich, Cat. No. 74255-250G), 10% glycerol (Carl Roth, Cat. No. 3783.1), 0.7% (v/v) 2-mercaptoethanol (Sigma, Cat. No. M3148-100ML), and bromophenol blue (Merck, Cat. No. 114405)) for 10 min at 95°C.

### Western Blot

Lysates (10 µL per lane) were separated on 10% SDS polyacrylamide gels in 1× SDS running buffer (25 mM Tris, 250 mM glycine (Glentham Life Sciences, Cat. No. GM2385), 3.45 mM SDS) alongside a protein ladder (GenScript, Cat. No. M00624-250). Proteins were transferred to a nitrocellulose membrane (NC Cytiva, Cat. No. 10600004) in 1× transfer buffer (25 mM Tris, 250 mM glycine, 20% ethanol (Chemsolute, Cat. No. 2212.5)). Membranes were blocked in 5% non-fat milk (Carl Roth, Cat. No. 145.2) in PBST (1× PBS (Carl Roth, Cat. No. 9150.1), 0.1% Tween-20 (Carl Roth, Cat. No. 9127.2)) for 1 h at RT and incubated with primary antibodies anti-CAPRIN1 (Abcam, Cat. No. ab244360; used at 0.2 µg/mL), anti-N (Invitrogen, Cat. No. MA5-36272; 0.67 µg/mL), anti-G3BP1 (Sigma, Cat. No. HPA004052; 0.4 µg/mL) and anti-β-actin (Proteintech, Cat. No. 66009-1-Ig; 0.1 µg/mL) overnight at 4°C. After three washes in PBST (5 min each), membranes were incubated with goat anti-rabbit IgG Alexa Fluor 647 (Invitrogen, Cat. No. A-21245; 0.4 µg/mL) and goat anti-mouse IgG Alexa Fluor 568 (Invitrogen, Cat. No. A-11004; 0.4 µg/mL) secondary antibodies in PBST for 1.5 h at RT, washed three times in PBST, and imaged with a VILBER Fusion FX7 EDGE V0.70 (Evolution-Capt Edge software).

### Infections for microscopy

One day before infection, cells were seeded to reach approximately 70% confluency on the day of infection. A549-ACE2-TMPRSS2 and CAPRIN1 KO cells were seeded at 5×10^4^ cells/well in 24-well plates (on 12 mm coverslips (Paul-Marienfeld, Cat. No. 112520)) or 3×10^4^ cells/well in 8-well µ-slides (ibidi, Cat. No. 80807); Vero E6 cells were seeded at 4×10^4^ cells/well in 24-well plates (on 12 mm coverslips) or 1×10^4^ cells/well in ibidi 8-well µ-slides. Cells were washed once with PBS and inoculated with the indicated virus in serum-free DMEM at the specified MOI. Plates were incubated for 1 h at 37°C with gentle agitation every 10 min. The inoculum was then removed and replaced with complete medium. The infected cells were maintained under standard conditions (37°C, 5% CO₂) for the duration of the experiment. Cells were fixed with 4.5% formaldehyde for 30 min at RT.

### Immunofluorescence (IF) staining of fixed cells

After fixation, cells were permeabilized with 0.1% Triton X-100 (Sigma-Aldrich, Cat. No. T8787-50ML) in PBS for 15 min and blocked with 3% (w/v) bovine serum albumin (BSA) (AppliChem/ITW Reagents, Cat. No. A6588.0100) in PBS for 5 min at RT. Cells were incubated with primary antibodies (Supplemental Table 1) at the indicated concentrations in PBS for 1 h at RT on a rocker, afterwards washed three times with 0.1% Tween-20 (Carl Roth, Cat. No. 9127.2) in PBS and then incubated with the corresponding secondary antibodies (Supplemental Table 2) at the indicated concentrations for 1 h at RT on a rocker. Where indicated, Hoechst 33342 (Merck, Cat. No. 14533-100MG) was included in the secondary antibody solution. Cells were washed three times with PBST. For N-hydroxysuccinimidyl-ester (NHS) staining, dyes (Thermo Fisher Scientific; catalog numbers listed in Supplemental Table 3) were applied after secondary antibody incubation and rocked for 1.5 h at RT. Samples were washed and stored at 4°C in the dark until imaging. When cells were seeded on coverslips, coverslips were washed once with ddH₂O and mounted cell-side down onto glass slides using 5 µL of ProLong Diamond Antifade Mountant (Invitrogen, Cat. No. P36961). Samples were dried overnight at RT in the dark.

**Supplemental Table 1:** Primary antibodies used for IF and U-ExM.

| Antibody | Stock Concentration | Host | Cat. No. | Company | Concentration IF | Concentration ExM |
| --- | --- | --- | --- | --- | --- | --- |
| N protein | 1 mg/mL | mouse IgG2b, mAB | A02048-100 | Genscript | 1 µg/mL | 2 µg/mL |
| dsRNA | 1 mg/mL | mouse IgG2a, mAB | Ab01299-2.0 | Absolute Antibody | 2 µg/mL | 5 µg/mL |
| G3BP1 | 0.1 mg/mL | rabbit IgG, pAB | HPA004052 | Atlas Antibodies | 0.5 µg/mL | 4 µg/mL |
| CAPRIN1 | 0.1 mg/mL | rabbit IgG, pAB | ab244360 | Abcam | 0.5 µg/mL | 4 µg/mL |
| CAPRIN1 | 0.1 mg/mL | rabbit IgG, pAB | HPA018126 | Atlas Antibodies | 0.5 µg/mL | 4 µg/mL |
| Calnexin | 0.9 mg/mL | rabbit IgG, pAB | ab22595 | Abcam | 0.9 µg/mL | 4.5 µg/mL |
| LAMP1 | 0.9 mg/mL | rabbit IgG, pAB | ab24170 | Abcam | 0.9 µg/mL | no specific signal |
| LAMP1 (clone H4A3) | 0.1 mg/mL | mouse IgG1 kappa, mAB | ab25630 | Abcam | 2 µg/mL | no specific signal |
| S protein (clone 1A9) | 1 mg/mL | mouse IgG1 kappa, mAB | ab273433 | Abcam | 2 µg/mL | no specific signal |
| Ultra-LEAF™ Purified CD63 | 1 mg/mL | mouse IgG1 kappa, mAB | 353040 | BioLegend | 2 µg/mL | 4 µg/mL |

**Supplemental Table 2:** Secondary antibodies used for IF and U-ExM.

| Antibody | Stock Concentration | Host | Cat. No. | Company | Concentration IF | Concentration ExM |
| --- | --- | --- | --- | --- | --- | --- |
| Alexa Fluor 405 Plus | 2 mg/mL | goat, anti-rabbit IgG (H+L) | A48254 | ThermoFisher Scientific | 2 µg/mL | 4 µg/mL |
| Alexa Fluor 488 | 2 mg/mL | Goat anti-Rabbit IgG (H+L) | A11008 | ThermoFisher Scientific | 2 µg/mL | 4 µg/mL |
| Alexa Fluor 488 | 2 mg/mL | Goat anti-Mouse IgG1 | A-21121 | ThermoFisher Scientific | 2 µg/mL | 4 µg/mL |
| Alexa Fluor 568 | 2 mg/mL | goat, anti-Mouse IgG2b | A21144 | ThermoFisher Scientific | 2 µg/mL | 4 µg/mL |
| Alexa Fluor 568 | 2 mg/mL | Goat anti-Mouse IgG (H+L) | A-11004 | ThermoFisher Scientific | 2 µg/mL | 4 µg/mL |
| Alexa Fluor 647 | 2 mg/mL | goat, anti-Mouse IgG2a | A21241 | ThermoFisher Scientific | 2 µg/mL | 4 µg/mL |

**Supplemental Table 3:** Dyes used for IF and U-ExM.

| Dye | Stock Concentration | Cat. No. | Company | Concentration IF | Concentration ExM |
| --- | --- | --- | --- | --- | --- |
| Hoechst 33342 | 10 mg/mL | 14533-100MG | Merck | 1 µg/mL | 10 µg/mL |
| NHS Ester + AF488 | 1 mg/mL | A20000 | ThermoFisher Scientific | 10 µg/mL | 10 µg/mL |
| NHS Ester + AF568 | 1 mg/mL | A20003 | ThermoFisher Scientific | 10 µg/mL | 10 µg/mL |
| NHS Ester + AF647 | 1 mg/mL | A20006 | ThermoFisher Scientific | 10 µg/mL | 10 µg/mL |

### Ultrastructure Expansion Microscopy (U-ExM)

U-ExM was performed on fixed cells based on a previously published protocol^28^ with modifications as detailed below.

Cells were seeded on 12 mm glass coverslips (Paul-Marienfeld, Cat. No. 112520) the night before infection and infected the next day as described above. Cells were incubated overnight at 37°C in anchoring solution containing 0.7% (v/v) formaldehyde (Sigma-Aldrich, Cat. No. 252549-1L) and 1% (v/v) acrylamide (Sigma-Aldrich, Cat. No. A4058-100ML) in PBS. A monomer solution containing 19% (w/v) sodium acrylate (Sigma-Aldrich, Cat. No. 408220-25G), 0.1% (v/v) N,N′-methylenebisacrylamide (Sigma-Aldrich, Cat. No. M1533-25ML), and 10% (v/v) acrylamide in PBS was prepared the day before and stored at -20°C.

On the following day, 10% (w/v) ammonium persulfate (APS; Sigma-Aldrich, Cat. No. 09913-100G) and 10% (v/v) N,N,N′,N′-tetramethylethylenediamine (TEMED; Sigma-Aldrich, Cat. No. T22500-100ML) were thawed on ice. A gelation chamber was assembled on ice using a parafilm-wrapped glass slide and a Petri dish lined with moist tissue to maintain humidity. Each coverslip was washed in PBS and excess liquid removed. A 45 µL gelation mix (35 µL monomer solution, 5 µL TEMED, 5 µL APS) was dispensed as a droplet onto the slide, and the coverslip was placed cell-side down onto the droplet. Gelation proceeded for 1 h at 37°C in the humid chamber.

Coverslips with attached gels were transferred to a 6-well plate containing denaturation buffer (200 mM SDS, 200 mM NaCl (Sigma-Aldrich, Cat. No. S5886-500G), 50 mM Tris pH 9) and incubated at RT on a rocker for ∼20 min until the gel detached from the coverslip. Gels were then transferred to 1.5 mL denaturation buffer in a 2 mL tube and denatured for 90 min at 95°C, or at 70°C when preservation of dsRNA was required. Gels were washed twice in ddH₂O for 15 min on a rocker and stored in ddH₂O at 4°C or washed in 50% (v/v) glycerol/ddH₂O and stored at -20°C for longer storage.

For immunostaining, gels were trimmed (using the lid of a 15 mL conical tube as a cutter), shrunk in PBS for 15 min, and blocked in 3% (w/v) BSA for 30 min on a shaker. Primary and secondary antibodies (Supplemental Table 1 and Supplemental Table 2) were incubated either for 2.5 h or overnight, depending on the antigen (Supplemental Table 4), and shaken on a rocker. Between primary and secondary antibody incubations, gels were washed three times in PBS 0.5% Tween-20 for 10 min each.

Scale bars for U-ExM images are reported as post-correction, using the expansion factor of 4.25× as determined by Liffner *et al.* (2023)^28^. They therefore refer to pre-expansion dimensions. The same factor was applied to samples denatured at 70°C, for which gel dimensions were comparable to those denatured at 95°C.

For imaging, poly-L-lysine (1:5 in PBS) (Sigma-Aldrich, Cat. No. P4707-50mL) was applied to a glass-bottom imaging dish (ibidi, Cat. No. 81158) for at least 5 min, then washed three times with ddH₂O. The gels were drained of excess water and were mounted on the dishes.

**Supplemental Table 4:** Incubation times of primary and secondary antibodies for U-ExM.

| Primary Antibody | Cat. No. | Incubation for<br>2.5 h at 37°C | Incubation<br>overnight at RT |
| --- | --- | --- | --- |
| N protein | A02048-100 | x |  |
| dsRNA | Ab01299-2.0 | x |  |
| G3BP1 | HPA004052 |  | x |
| CAPRIN1 | ab244360 |  | x |
| CAPRIN1 | HPA018126 | x |  |
| Calnexin | ab22595 | x |  |
| Ultra-LEAF™<br>Purified CD63 | 353040 |  | x |
| Secondary<br>Antibody | Cat. No. | Incubation for<br>2.5 h at RT | Incubation<br>overnight at RT |
| Alexa Fluor 405 Plus | A48254 |  | x |
| Alexa Fluor 488 | A11008 | x |  |
| Alexa Fluor 488 | A-21121 | x |  |
| Alexa Fluor 568 | A21144 | x |  |
| Alexa Fluor 568 | A-11004 | x |  |
| Alexa Fluor 647 | A21241 |  | x |

### Imaging of samples

Fixed samples were imaged on a Nikon Ti2 microscope equipped with a Yokogawa CSU-W1 spinning-disc confocal unit, Mad City Labs Nano-Drive piezo system, and a Hamamatsu ORCA-Fusion BT sCMOS camera and operated with NIS-Elements AR 6.10.01. Immunofluorescence samples were imaged in spinning-disc confocal mode using a 100× oil objective (NA 1.49). Expanded (U-ExM) samples were imaged using a 60× water objective (NA 1.27). Fluorophores were excited with 405/488/561/638 nm laser lines and a quad dichroic (405/488/568/647 nm), and emission was collected through a quad bandpass filter (446/510/581/703 nm). Z-stacks were acquired with a step size of 0.2 µm. Displayed images are single confocal planes.

### Processing of images

All images were processed using Fiji. Processing was limited to pseudo-colouring and adjusting brightness and contrast for display purposes. This was applied uniformly across the entire image and equally to all compared conditions. Images were cropped to the same dimensions within each experiment so that the scale bar applies to all images. Morphological scoring (Figure 9) was performed manually on all fields of view. Only N-positive cells were scored, and each phenotype was scored independently.

### Generation of CAPRIN1 KO cells

CAPRIN1-knockout A549 cells expressing ACE2 and TMPRSS2 were generated by CRISPR/Cas9-mediated gene editing to introduce frameshift mutations in the CAPRIN1 coding sequence^29^. Two guide RNAs (gRNAs) targeting the first translated exon were designed using Benchling and CHOPCHOP, selecting for predicted high on-target and low off-target scores. The selected sgRNA sequences were 5’-GGCCATGAAGCAGATTCTCG-3’ and 5’-CGACAAGAAACTTCGGAACC-3’.

Complementary oligonucleotides with BbsI-compatible overhangs were annealed and cloned into the BbsI-digested (NEB, Cat. No. R3539S) pSpCas9(BB)-2A-Puro (PX459) V2.0 vector (Addgene, Cat. No. 62988). Sequence verified constructs (pSpCas9-CAPRIN1-gRNA1 and pSpCas9-CAPRIN1-gRNA2) were transfected into A549-ACE2/TMPRSS2 cells using Lipofectamine 2000 (Invitrogen, Cat. No. 11668030) according to the manufacturer’s instructions. Briefly, 3.5×10^5^ cells were seeded into a 6-well to reach ∼80% confluency the following day. Per well, 2.5 µg total DNA (2.25 µg CRISPR constructs + 0.25 µg pcDNA3-mNeonGreen^30^) was complexed with 9 µL Lipofectamine 2000 in Opti-MEM (Gibco, Cat. No. 31985-062). Complexes were incubated with cells for 6 h, after which the medium was replaced with complete DMEM/F-12 supplemented with 10% FBS, 0.5 µg/mL puromycin, and 10 µg/mL blasticidin.

To enrich for transfected cells, single cells were sorted by fluorescence-activated cell sorting (FACS). Sorting was performed on a BD FACS Aria Fusion equipped with a 50 mW 488 nm laser and emission filter 525/50. The nozzle size was 70 µm and BD FACS Diva Software v.8.0.1 was used. SSC and FSC measurements were performed with the 488 nm laser. Gating selected the brightest 3% of mNeonGreen-positive cells and sorted them individually into a 96-well plate containing complete media. Single cell clones were expanded.

For genotyping, cells were lysed directly in 96-well plates using DirectPCR Cell Lysis Reagent (VWR, Cat. No. 302-C) supplemented with proteinase K (Thermo Fisher Scientific, Cat. No. EO0491; 55°C for 6 h, then heat-inactivated at 85°C for 45 min). Crude lysates served as templates for PCR amplification of the targeted genomic region (forward primer 5’-CTTCTCTCTCCTTGCGGTCTG-3’ and reverse primer 5’-CGCTCCCGAGACGAAAAGAA-3’) using Q5 DNA polymerase (NEB, Cat. No. M0491S). PCR products were purified using the QIAquick PCR Purification Kit (Qiagen, Cat. No. 28106). As a first screen, the PCR products were Sanger sequenced in bulk, and the chromatograms were analyzed for insertions and deletions using TIDE^31^. Clones that likely contained multiple genotypes were followed up by cloning the PCR products into a vector with CloneJet (Thermo Fisher Scientific, Cat. No. K1231) to sequence individual genotypes. At least eight plasmids per single cell clone were sequenced, and only if three genotypes were identified that all contained a frameshift was the clone considered a putative KO.

Putative KO clones were validated via IF and WB. Immunofluorescence staining with an anti-CAPRIN1 antibody (Abcam, Cat. No. ab244360) tested for loss of cytoplasmic CAPRIN1 signal. Western blot analysis tested for loss of CAPRIN1 protein in total cell lysate. Approximately 1×10^6^ cells in a 6-well were washed twice with cold PBS, scraped into 1 mL PBS per well, and pelleted by centrifugation (3,000 × g, 5 min, 4°C). Pellets were resuspended in 120 µL of 1× SDS loading buffer (50 mM Tris pH 6.8, 2% SDS, 10% glycerol, 0.7% (v/v) 2-mercaptoethanol, and bromophenol blue) and denatured at 95°C for 10 min. Verified KO cell lines were cryopreserved.

### Statistics and visualization

Graphs were generated in Python using the matplotlib library. The Fisher’s exact test and Mann-Whitney U test were performed with the Python scipy library.

## Results

### Meta-analysis of N protein interactomes identifies CAPRIN1 as an SG protein of interest

To identify candidate host proteins that interact with the SARS-CoV-2 N protein, we first performed a meta-analysis of 17 published N interactome screens from 14 studies^12,32–44^ (Supplemental Table 5). All of these studies involved exogenous N expression outside of the context of viral infection. We observed low overlap of individual proteins across datasets, likely reflecting differences in cell type, interaction systems, sample preparation protocols, and data processing pipelines (Figure 1A). To overcome this heterogeneity, we mapped all identified host proteins onto a human protein-protein interaction network (PICKLE human interactome, release 3.3^27^) and performed sub-clustering analysis. This revealed a subcluster that contained SG proteins (Figure 1B), such as the well-characterized proteins PABPC1^45^ and UBAP2L^46^, consistent with the reports from other studies^14,36^.

**Figure 1:**
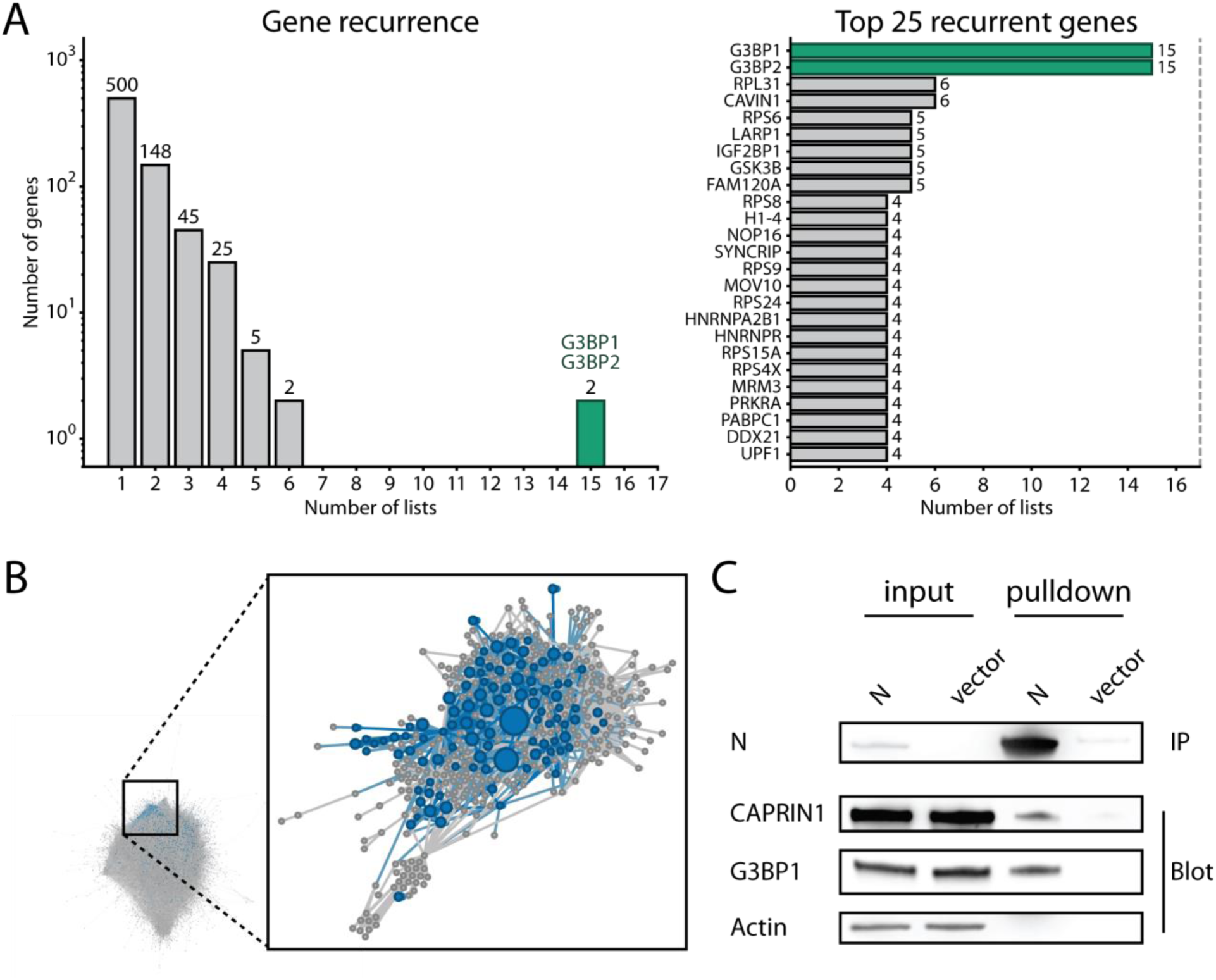
CAPRIN1 is a stress granule protein that binds to N. (A) Recurrence plots illustrating the overlap of N interactors from 17 mass spectrometry screens. The proteins with the highest recurrence are highlighted in green. (B) Mapping of published N interactors (blue) onto the PICKLE human interactome, release 3.3 (grey). The zoom-in shows the Louvain clustering that contains stress granule proteins. Each node represents a single protein, and the diameter of the node is proportionate to the number of MS screens it was identified in. (C) Lenti-X 293T cells were transfected with an empty or N expression vector. Co-immunoprecipitation against N was followed by Western blot against the N interactors CAPRIN1 and G3BP1 and the negative IP control actin.

G3BP1 has the greatest consensus between the datasets, appearing in 15 (Figure 1A), and is the most extensively characterized N protein interactor, with its binding interface resolved at atomic resolution and biological relevance confirmed both *in cellulo* and *in vivo*^15,20^. Therefore, we chose G3BP1 as the reference protein in this study. CAPRIN1 was selected based on its central role in promoting SG condensation through its direct interaction with G3BP1^21,22^, its contribution to PKR-dependent innate immune signalling within SGs^17^, and the absence of spatial characterization in SARS-CoV-2 infection.

To test whether CAPRIN1 associates with the N protein, Lenti-X 293T cells were transfected with an expression construct for the N protein, followed by co-immunoprecipitation and Western blot (WB) analysis. Our findings showed that CAPRIN1 co-precipitated with the N protein along with G3BP1 but not the actin negative control (Figure 1C).

**Supplemental Table 5:** Methodologies of N interactome studies.

| First Author | Cell Type | System |
| --- | --- | --- |
| Armstrong SD. et al. | HEK293T | AP-MS eGFP |
| Chen Z. et al. | HEK293T | proximity labelling BioID2 |
| Gordon DE. et al. | HEK-293T/17 | AP-MS StrepII |
| Kim DK. et al. | yeast | Yeast two-hybrid |
| Laurent EMN. et al. | HEK293T | proximity labelling BirA |
| Li J. et al. | yeast | Yeast two-hybrid |
| Liu X. et al. | HEK293 | AP-MS StrepII |
| Liu X. et al. | HEK293 | proximity labelling BirA |
| May DG. et al. | A549 | proximity labelling BioID2 |
| Min YQ. et al. | 293T | AP-MS S-tag |
| Nabeel-Shah S. et al. | HEK293 | AP-MS eGFP |
| Samavarchi-Tehrani P. et al. | A549 | proximity labelling miniTurbo |
| Stukalov A. et al. | A549 | AP-MS HA-tag |
| Zheng X. et al. | HEK293T | AP-MS StrepII |
| Zheng X. et al. | Calu-3 | AP-MS StrepII |
| Zhou Y. et al. | yeast | Yeast two-hybrid |
| Zhou Y. et al. | Caco-2 | AP-MS FLAG |

### Staging of SARS-CoV-2 infections identifies LAMP1-positive membrane-bound egress compartments with intraluminal content

To determine where G3BP1 and CAPRIN1 are localized in the SARS-CoV-2 viral cycle, we first defined the distribution of viral proteins at replication and egress sites. Building on the staging framework established by Scherer *et al.* (2022)^47^, we distinguished two stages of the viral cycle by the sequential appearance of the N and S proteins: N without S marking replication and N together with S marking egress (Figure 2A). The N protein is the first structural protein to be detected in infected cells and therefore can serve as an early marker for viral replication^47^. At this stage, only the N protein is detectable and appears in small clusters surrounding the double-stranded RNA (dsRNA), indicating active viral replication in DMVs derived from the ER. The absence of S protein during this phase is consistent with the temporal hierarchy of structural protein expression, in which N protein production precedes that of the transmembrane structural proteins S, M, and E^47^. As the infection progresses, the growth of DMVs results in a decrease in the calnexin signal, creating a void-like area in the ER (Figure 2B), which is consistent with the reorganization of the ER into DMVs observed in other microscopy studies^47,48^.

**Figure 2:**
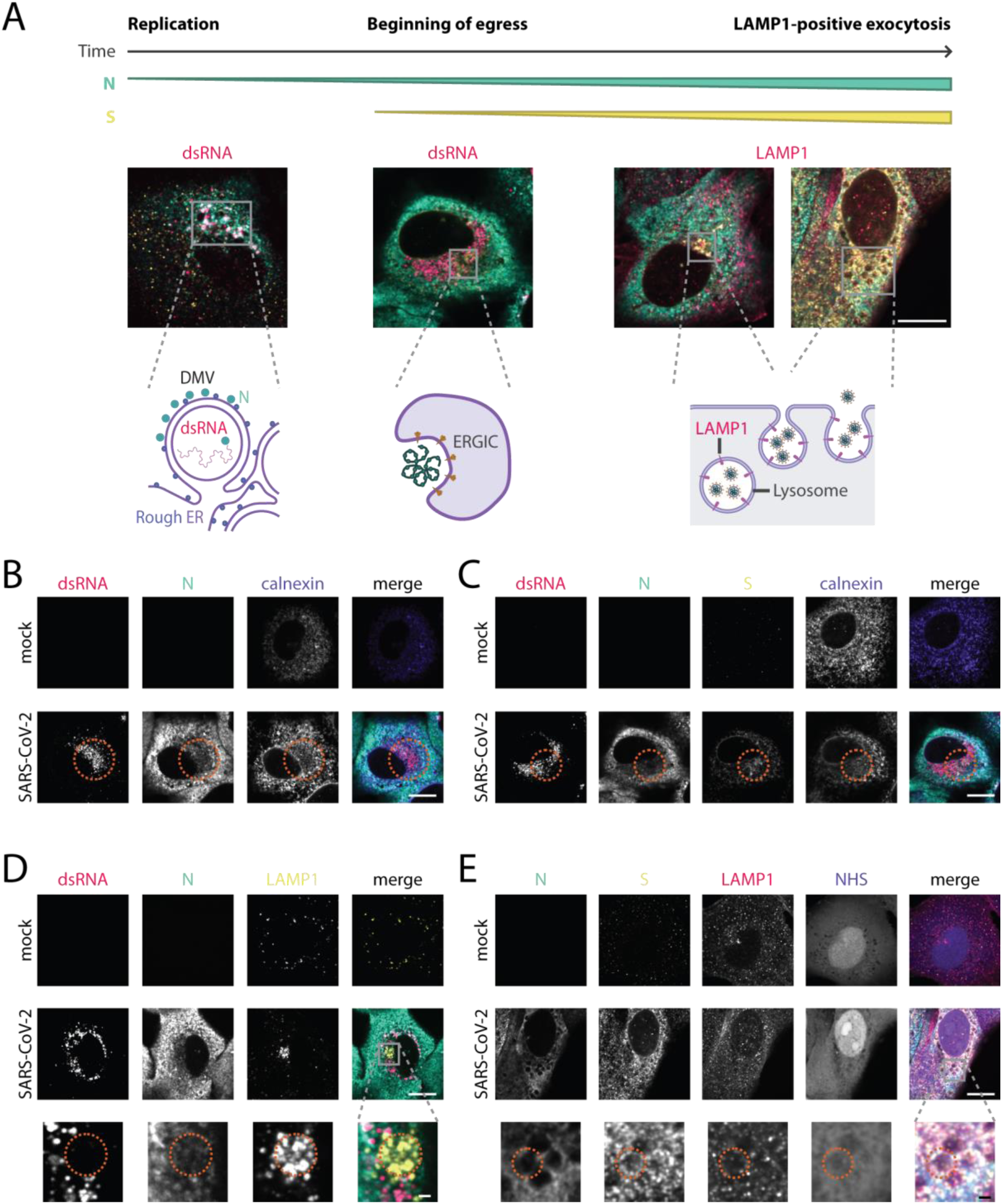
Identification of SARS-CoV-2 replication and egress compartments relative to cellular markers. A549 cells were infected with SARS-CoV-2 at an MOI of 5 and stained by IF for the indicated proteins. Scale bars, 10 µm; Zoom-in scale bars, 1 µm. (A) Schematic illustrating the progression of viral infection. N (green) with dsRNA (red) marks early replication sites, whereas N with S (yellow) marks egress-associated, LAMP1 (red)-positive compartments. Cells were fixed at 6 hpi or 11 hpi. (B) ER is reorganized due to growth of DMVs, leaving a region of reduced calnexin signal in perinuclear ER of infected cells (red circle). Cells were fixed at 11 hpi and stained for dsRNA (red), N (green), and the ER marker calnexin (purple). (C) S (yellow) appears spatially separated from dsRNA (red) hubs, consistent with S protein marking the location of nascent egress sites (red circle). Cells were additionally stained for N (green) and calnexin (purple) and fixed at 11 hpi. (D) As infection progresses, LAMP1-positive compartments are redistributed to the perinuclear region (red circle) rather than remaining evenly distributed throughout the cell. Cells were fixed at 11 hpi and stained for dsRNA (red), N (green), and LAMP1 (yellow). (E) Colocalization of S protein (yellow) and LAMP1 (red) marks the LAMP1-positive egress-associated compartments (red circle): the two signals first appear together in the perinuclear region and subsequently colocalize on membrane-rich organelles containing intraluminal content visualized in the NHS channel (purple). Cells were additionally stained for N (green) and fixed at 11 hpi.

At the egress sites, S protein was detectable in addition to the N protein, partially or fully separated from the dsRNA-positive replication region (Figure 2C). This spatial separation is consistent with a model in which newly synthesized viral RNA is exported from DMVs and encapsulated by N protein on the cytoplasmic face of the replication organelles to form RNP complexes, which relocate to ERGIC-derived membranes where S, M, and E proteins are already positioned for virion assembly^4,49^. Based on these observations, S protein has been established as a SARS-CoV-2 egress marker^47^. Consistent with previous work^50^, LAMP1 was concentrated in the perinuclear region (Figure 2D) and colocalized with S protein on distinct compartments (Figure 2E). These resemble the LAMP1-positive organelles reported to undergo exocytosis during β-coronavirus egress^5,7,47^.

To further characterize the egress compartments, we stained cells with a fluorescent N-hydroxysuccinimidyl-ester (NHS), a chemical that labels amines and therefore provides a total protein reference background stain in cells. NHS staining revealed large 1-2 µm in diameter membrane-bound compartments with intraluminal content that were positive for N, S, and LAMP1 on their limiting membranes (Figure 2E). Collectively, these observations reproduce the published staging framework in our system and identify a membrane-bound compartment with intraluminal content as the structure to be characterized further.

### CD63-positive MVB-like compartments are involved in SARS-CoV-2 egress

Previous studies of SARS-CoV-2 have relied primarily on LAMP1 staining and conventional IF to identify egress compartments. However, LAMP1 is not exclusive to lysosomes but is present across compartments of the endolysosomal continuum^51^. In addition, conventional IF provides only limited resolution of whether a signal lies on the limiting membrane or in the lumen. To address both limitations, we combined U-ExM with staining for CD63, which is a marker used primarily to identify multivesicular bodies (MVBs) and localizes to the limiting membrane and to intraluminal vesicles (ILVs)^9,52^. U-ExM is a technique that physically enlarges biological samples and thereby increases the resolution of a conventional microscope approximately four-fold, enabling the visualization of subcellular structures that would otherwise remain unresolved^53,54^ (Figure 3A). By performing U-ExM on infected cells, we could show that the membrane-bound compartments (Figure 2E) clearly contain proteinaceous intraluminal content (Figure 3B).

**Figure 3:**
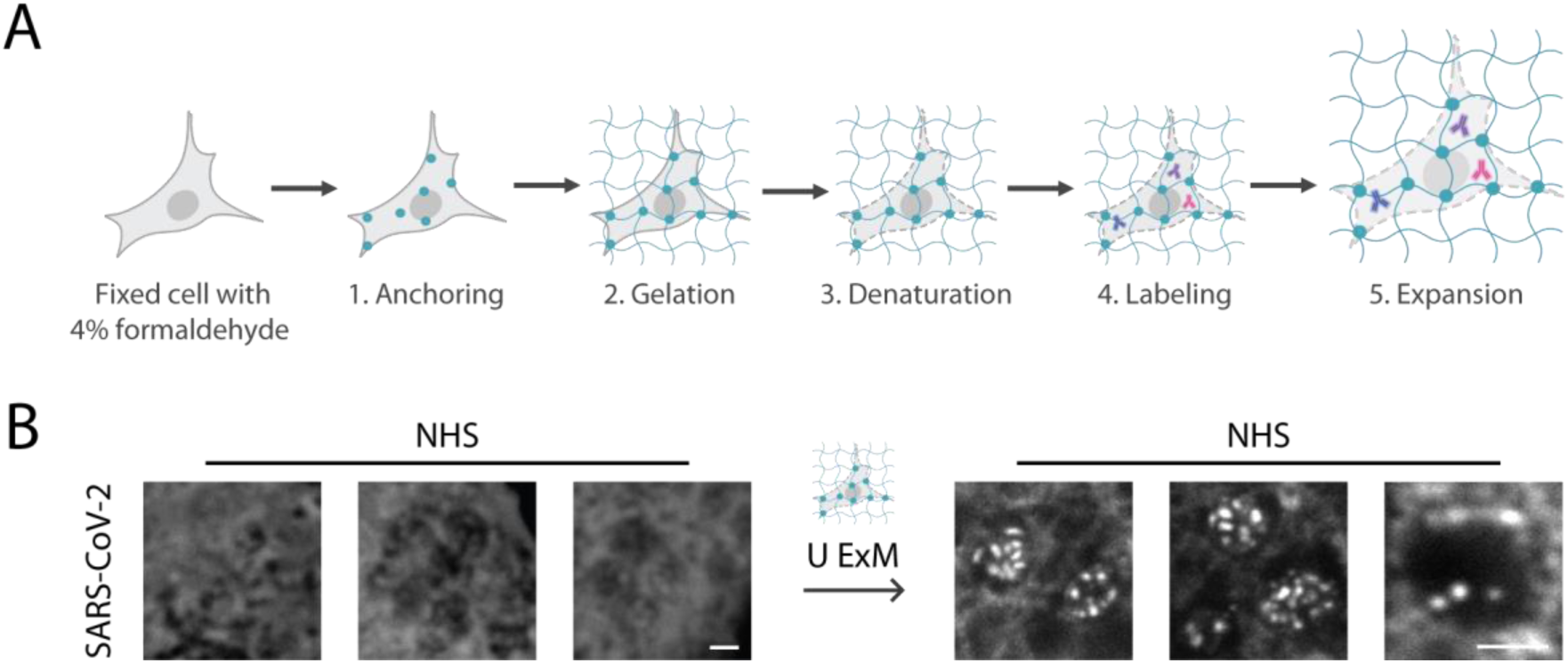
U-ExM generates higher spatial resolution images that highlight proteinaceous intraluminal content. (A) Schematic of the U-ExM protocol. 1: Anchoring: the fixed sample is incubated in formaldehyde/acrylamide to link proteins into the forming gel in the next step. 2: Gelation: the sample is embedded in a polyacrylamide/polyacrylate hydrogel. 3: Denaturation: heat- and SDS-mediated homogenization unfolds proteins and ensures expansion. 4: Labelling: incubation with primary antibody, secondary antibody, and additional dyes like NHS or Hoechst. 5: Expansion: immersion of the sample in water to physically expand it. (B) Zoom-in of putative egress compartments from IF and U-ExM processed SARS-CoV-2 infected cells stained with NHS (grey). A549 cells were infected with SARS-CoV-2 at an MOI of 5 and processed at 11 hpi. Zoom-in scale bars, 1 µm. For U-ExM images, the scale bars are post-correction. Each image is from an independent cell.

To further characterize the membrane-bound compartments (Figure 2E), we stained the samples for an additional marker. Staining for both LAMP1 and CD63 revealed that the LAMP1-positive compartments were also positive for CD63 (Figure 4A). By combining U-ExM with CD63 staining, we could show that these compartments contain CD63 on both the limiting membrane and within the lumen (Figure 4B). The luminal CD63 signal suggests that these compartments contain ILVs, consistent with the membranous structures previously observed inside SARS-CoV-2 egress compartments by TEM^8^. CD63 co-occurred with LAMP1 on the compartments, consistent with an MVB-like identity within the late-endolysosomal continuum^51^ (Figure 4A). Two independent LAMP1 antibodies showed the same localization to these compartments, supporting the specificity of the signal (Figure 4A and C). However, neither of the LAMP1 antibodies tested yielded specific signal in U-ExM samples. Therefore, LAMP1 was imaged by only conventional IF (Figure 4A and C), and its localization to ILVs remains unresolved. In U-ExM, these compartments contain N on the limiting membrane and in ILVs (Figure 4B), the latter likely containing other structural proteins such as S (Figure 2E), suggesting that these MVB-like compartments contain infectious virions. Together, we demonstrate that the SARS-CoV-2 egress compartments previously characterized as lysosomes based on LAMP1 staining alone are CD63-positive MVB-like compartments that contain N on the limiting membrane and in the lumen, suggesting that these compartments carry virions inside (Figure 4D).

**Figure 4:**
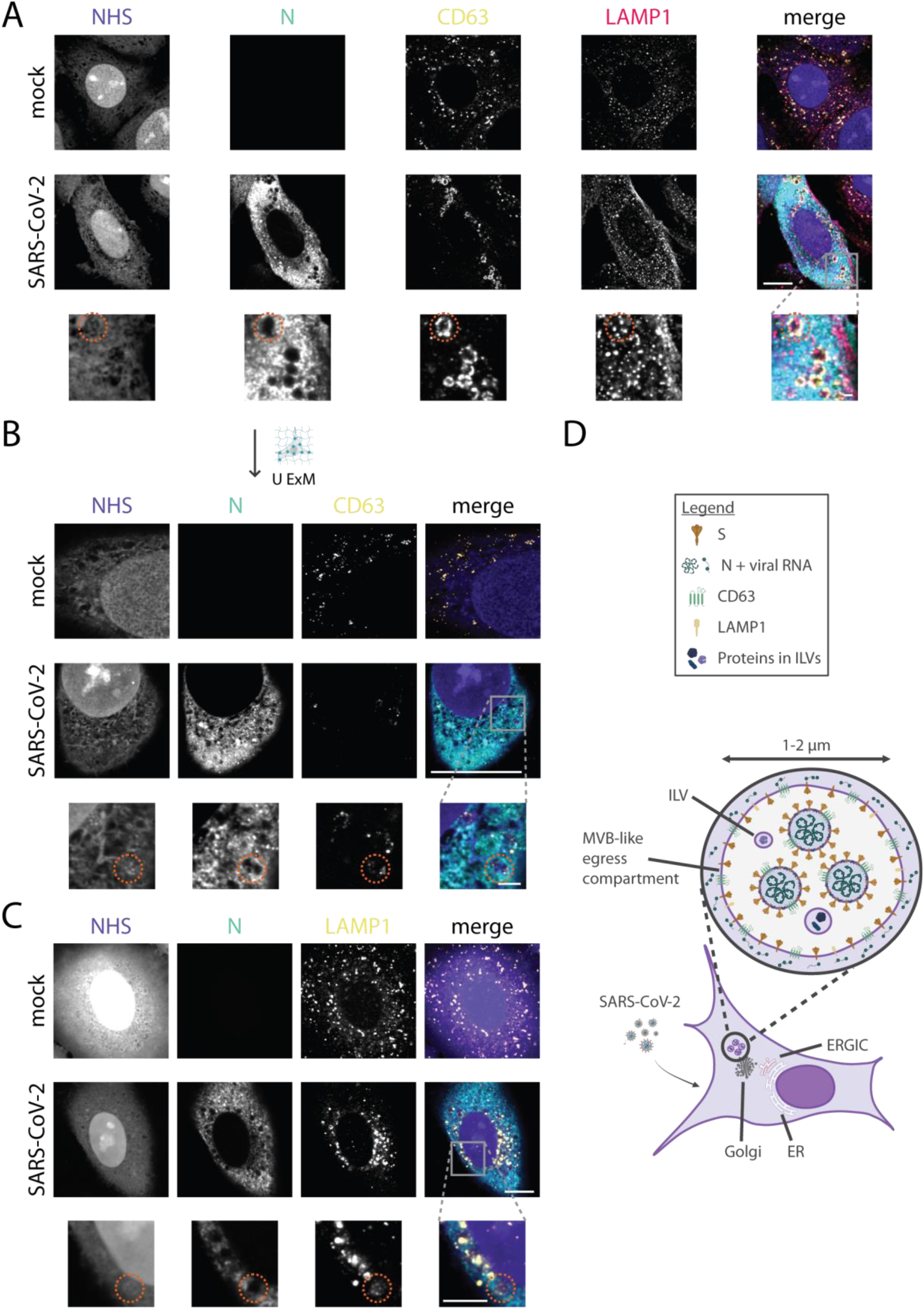
CD63 is also present on LAMP1-positive, MVB-like SARS-CoV-2 egress compartments. A549 cells were infected with SARS-CoV-2 at an MOI of 5 and processed at the indicated time points. Putative egress compartments are marked with a red circle. Scale bars, 10 µm; Zoom-in scale bars, 1 µm. For U-ExM images, the scale bars are post-correction. (A) Cells were fixed at 11 hpi and stained for NHS (purple), N (green), CD63 (yellow), and LAMP1 (red). (B) The cells were processed for U-ExM and stained for NHS (purple), N (green), and CD63 (yellow). (C) Cells were fixed at 6 hpi and stained with an alternate LAMP1 antibody (yellow) in addition to NHS (purple) and N (green). (D) Proposed model of the MVB-like compartment involved in SARS-CoV-2 egress. LAMP1 and CD63 reside on the limiting membrane together with S and N. CD63, as well as the viral S and N proteins, is also found in the intraluminal content, which likely represents virions. These compartments are 1-2 µm in diameter.

### G3BP1 but not CAPRIN1 is recruited to the periphery of DMVs

Having defined the replication and egress sites, we next investigated whether G3BP1 or CAPRIN1 is localized at the replication sites. To this end, we performed immunofluorescence staining with the combination of N protein, dsRNA, and the two SG proteins. In U-ExM, NHS was included as an additional reference to confirm the ultrastructural context of the replication area.

Interpreting protein colocalization at replication sites requires considering their topology. The dsRNA resides inside the DMV lumen, whereas SG proteins together with N localize to the cytoplasmic face of the DMV membrane. Although DMVs can reach diameters of up to 800 nm at late stages of infection^55^, conventional light microscopy cannot reliably separate luminal from cytoplasmic-facing signal. In U-ExM, the physical expansion resolves the DMV interior from its cytoplasmic exterior, and therefore the effectively increased resolution should spatially separate the luminal dsRNA from the cytoplasmic SG proteins.

Starting with the reference protein, G3BP1 was found to be enriched along with N around the dsRNA foci in the replication area (Figure 5A), consistent with the recruitment of G3BP1 by N to the replication site reported in other studies^18,56^. In U-ExM, the spatial separation between G3BP1 and dsRNA became apparent (Figure 5B), with G3BP1 localizing to the periphery of the dsRNA-positive structures, consistent with its expected position on the cytoplasmic face of the DMVs^57^.

**Figure 5:**
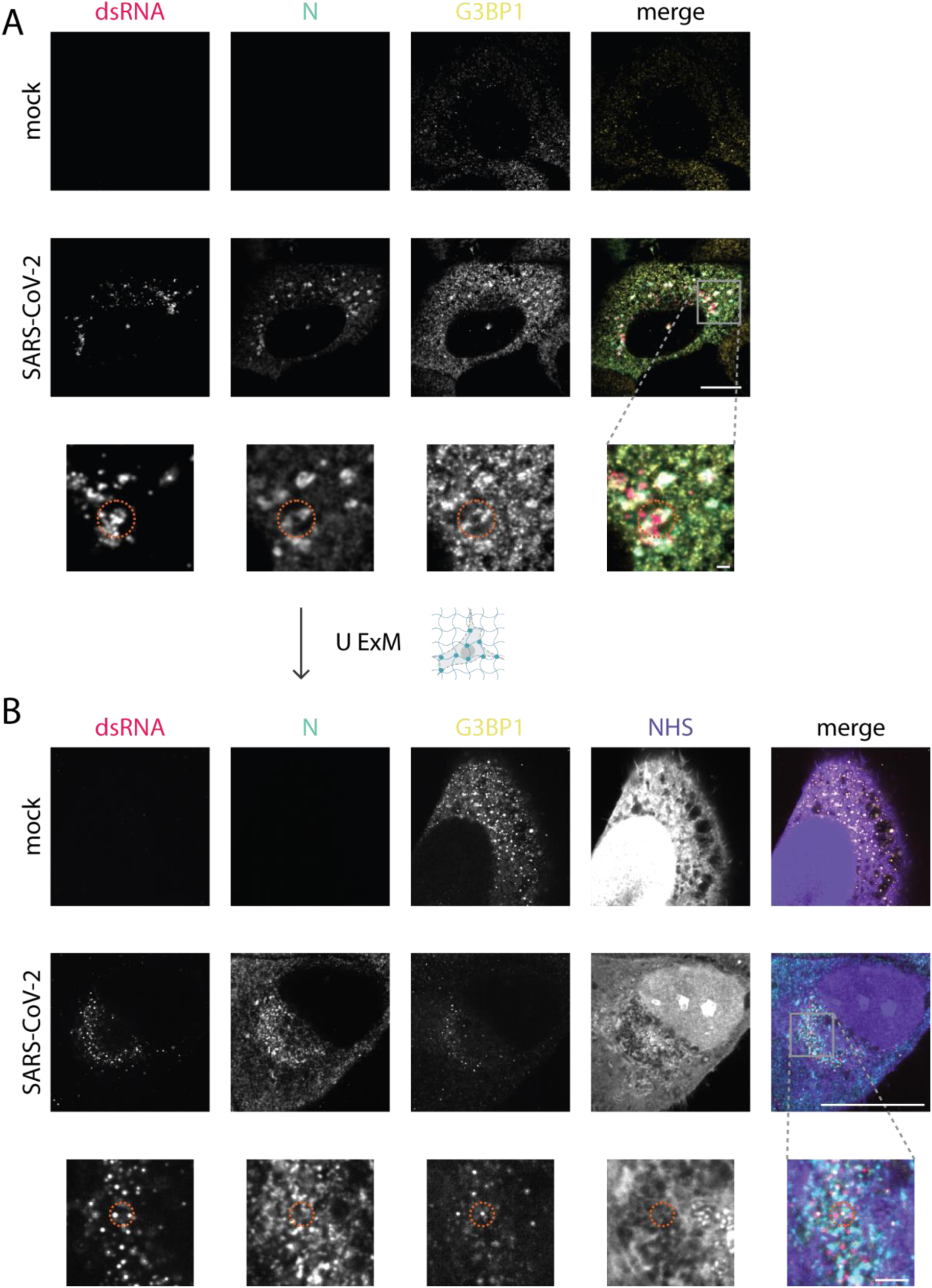
G3BP1 is recruited to SARS-CoV-2 replication sites. A549 cells were infected with SARS-CoV-2 at an MOI of 5 and processed for IF and U-ExM at the indicated times post infection. Replication sites are marked with a red circle. Scale bars, 10 µm; Zoom-in scale bars, 1 µm. For U-ExM images, the scale bars are post-correction. (A) Infected cells were fixed at 6 hpi and stained for G3BP1 (yellow), dsRNA (red), and N (green). (B) Cells at 11 hpi were processed for U-ExM and stained for G3BP1 (yellow), dsRNA (red), N (green), and NHS (purple). Colocalization of N and G3BP1 surrounding dsRNA was used to identify DMV replication sites.

CAPRIN1, by contrast, remained dispersed throughout the cytoplasm with no enrichment at replication sites, matching its distribution in uninfected cells (Figure 6). This is expected based on a model of competition between N and CAPRIN1 for G3BP1. A substantial pool of CAPRIN1 is already free of G3BP1 in unstressed cells^22^, and CAPRIN1 and N bind the same site on G3BP1^15,22,58^. Therefore, G3BP1 recruited via N is unavailable to CAPRIN1, leaving the free pool unchanged. Together, these results indicate that CAPRIN1 is not recruited to replication sites, in contrast to G3BP1. However as shown below, this did not mark the end of its involvement.

**Figure 6:**
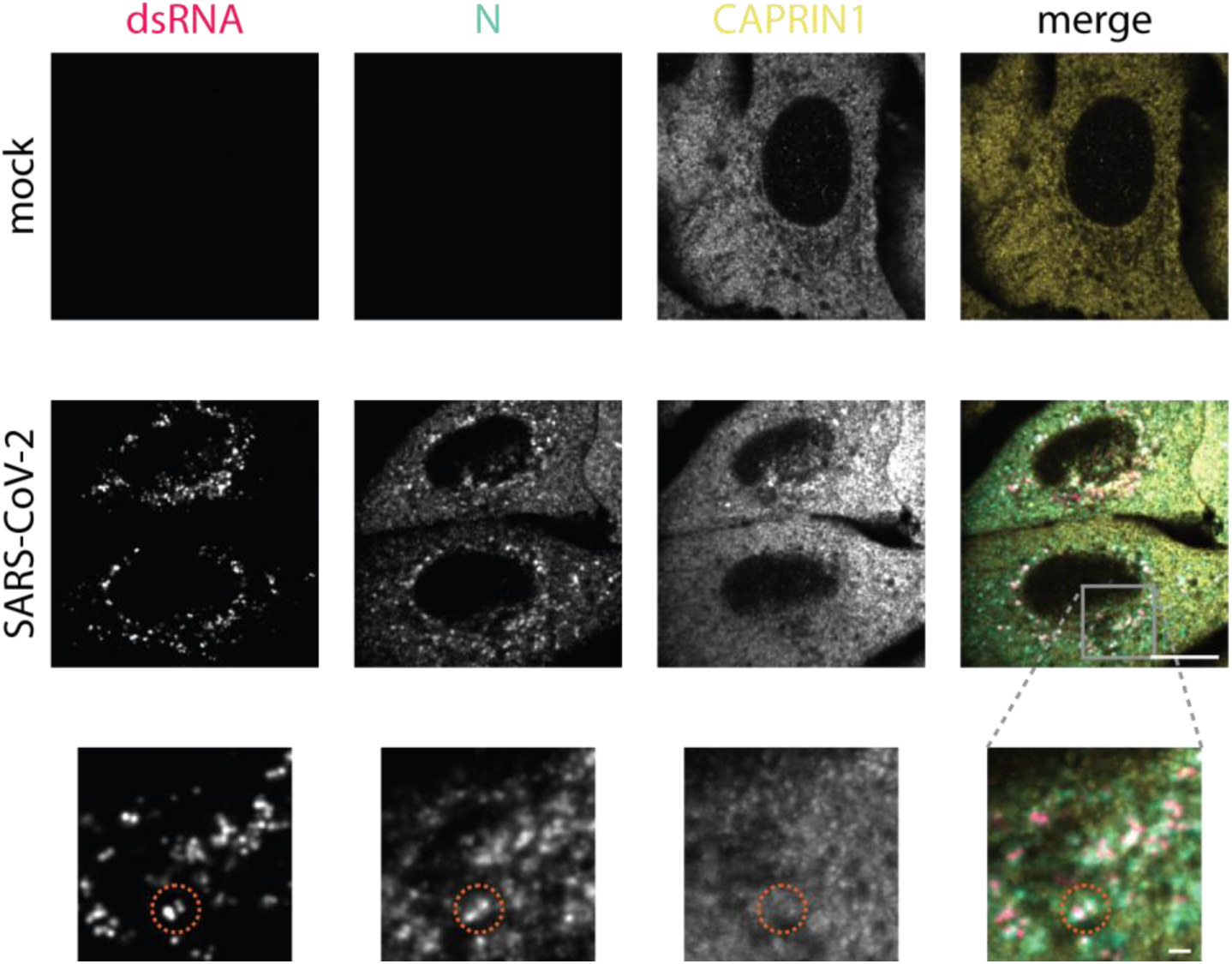
CAPRIN1 is not recruited to SARS-CoV-2 replication sites. A549 cells were infected with SARS-CoV-2 at an MOI of 5 and processed for IF at 6 hpi. Replication sites are marked with a red circle. Scale bars, 10 µm; Zoom-in scale bars, 1 µm. Infected cells were fixed and stained for CAPRIN1 (yellow), dsRNA (red), and N (green). Colocalization of N and dsRNA was used to identify DMV replication sites.

### U-ExM resolves distinct positions of G3BP1 and CAPRIN1 within the egress compartments

Since CAPRIN1 was not recruited to the replication sites, we investigated whether it is involved at a later stage of infection. To test this, two different staining combinations in IF were performed: 1) N protein with the two SG proteins and LAMP1, to evaluate co-occurrence with lysosomal markers, and 2) N protein with the two SG proteins and S, to evaluate co-occurrence with viral egress markers. In both staining combinations, we co-stained for dsRNA to identify egress sites by the lack of dsRNA. In U-ExM, we stained for the two SG proteins together with N and NHS to resolve the distribution of SG proteins relative to the MVB-like compartment ultrastructure and N. Because NHS labels amine groups on proteins but membranes are not strictly preserved during U-ExM, the position of the limiting membrane was inferred from the boundary of the NHS-dense signal. CD63 signal aligned with the outer edge of the NHS-dense region, supporting this assignment.

The increased resolution of U-ExM revealed a difference in phenotype between G3BP1 (Figure 7) and CAPRIN1 (Figure 8). In IF, G3BP1 and CAPRIN1 were both detected at ring-like structures positive for both S and LAMP1, indicating their presence at egress sites (Figure 7A and B, Figure 8A and B). However, U-ExM revealed that each protein occupied a distinct position within these compartments (Figure 7C, Figure 8C). G3BP1 localized predominantly to the limiting membrane of the MVB-like compartments, where it colocalized with N. In contrast, signal for CAPRIN1 was present on the limiting membrane of the MVB-like compartments and on the protein-dense material in the interior (additional MVBs are shown in Supplemental Figure 2). We also performed this experiment in Vero E6 cells, a cell line commonly used as a model for SARS-CoV-2 infection, and observed the same phenotype (Supplemental Figure 3). Together, these results show that two SG proteins that show a similar phenotype at egress sites by conventional IF occupy distinct positions within MVB-like compartments when resolved by U-ExM: G3BP1 primarily on the limiting membrane with N, and CAPRIN1 on both the limiting membrane and the ILVs.

**Figure 7:**
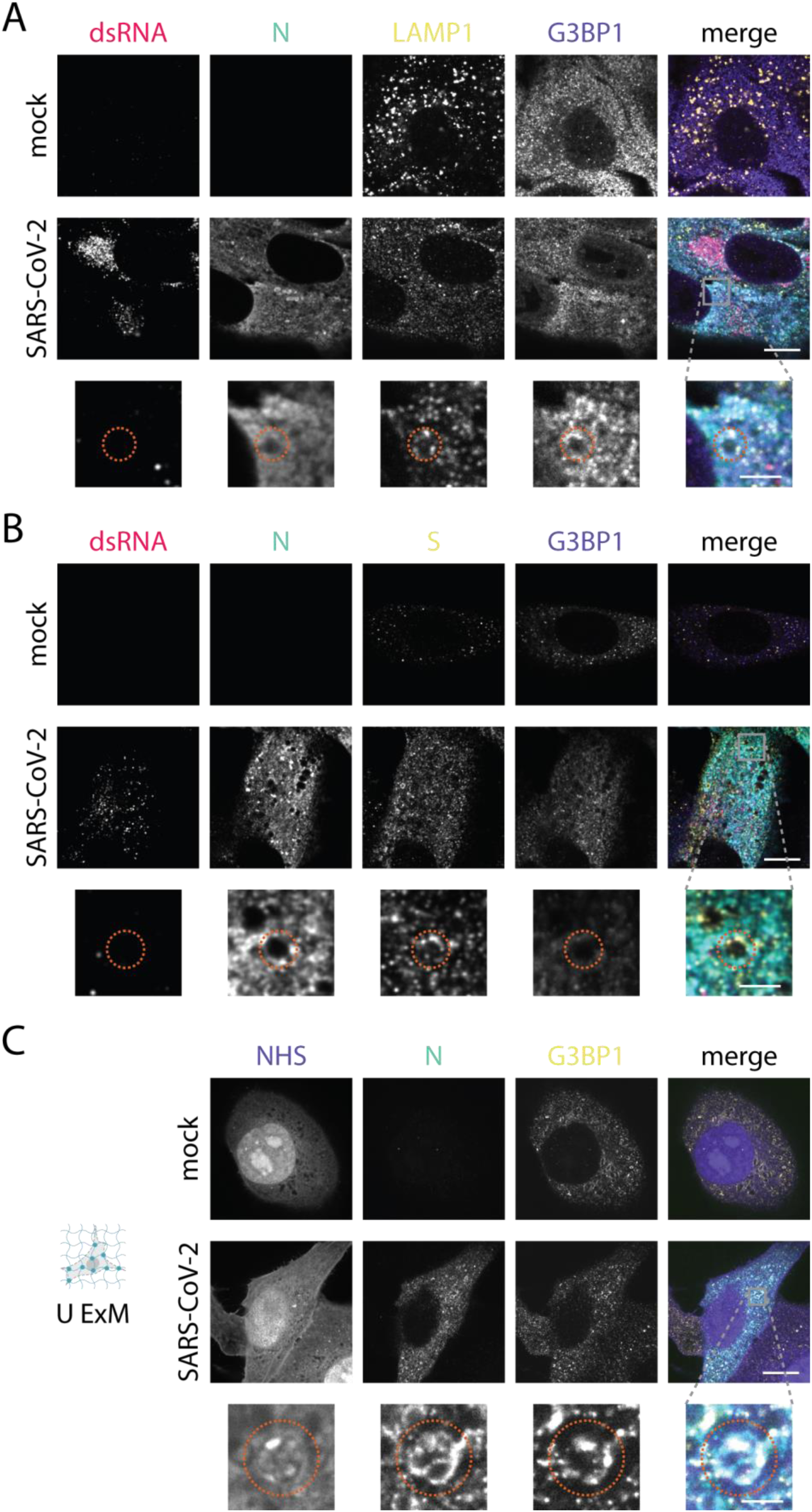
G3BP1 is present predominantly on the limiting membrane of MVB-like compartments. A549 cells were infected with SARS-CoV-2 at an MOI of 5 and processed for IF and U-ExM at the indicated times post-infection. The MVB-like compartments are marked with a red circle. Scale bars, 10 µm; Zoom-in scale bars, 1 µm. For U-ExM images, the scale bars are post-correction. (A) Infected cells were fixed and stained for G3BP1 (purple), dsRNA (red), N (green), and LAMP1 (yellow). The dsRNA was chosen as a counterstain to exclude the replication sites. Cells were fixed at 11 hpi. (B) Infected cells were fixed and stained for G3BP1 (purple), dsRNA (red), N (green), and S (yellow). Colocalization of N and S was used to identify egress compartments. Cells were fixed at 11 hpi. (C) Infected cells were processed for U-ExM and stained for G3BP1 (yellow), NHS (purple), and N (green). NHS was used to identify MVB-like compartments. Cells were fixed at 6 hpi.

**Figure 8:**
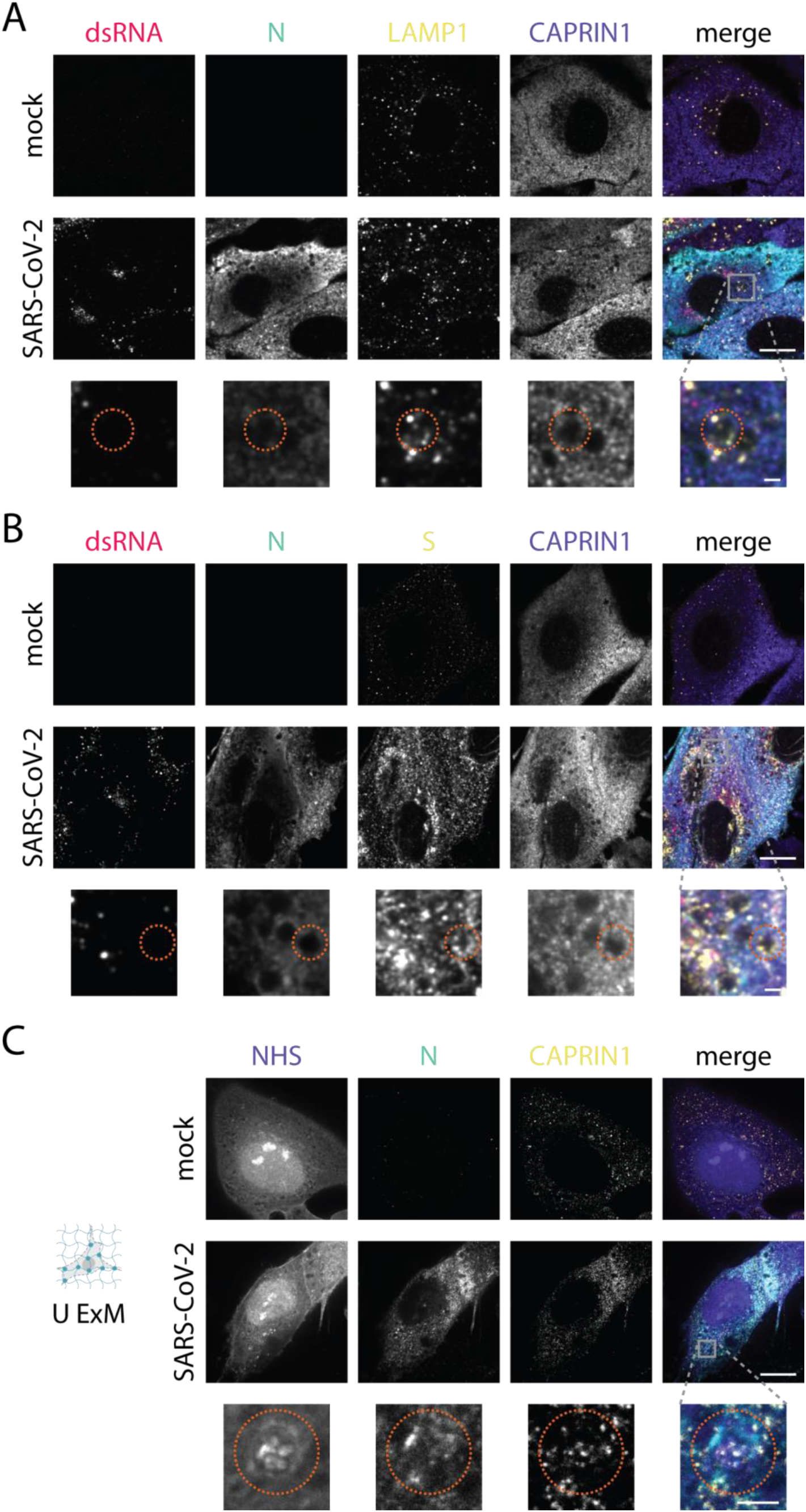
CAPRIN1 is present on the limiting membrane and on the intraluminal content of MVB-like compartments. A549 cells were infected with SARS-CoV-2 at an MOI of 5 and processed for IF and U-ExM at the indicated times post-infection. The MVB-like compartments are marked with a red circle. Scale bars, 10 µm; Zoom-in scale bars, 1 µm. For U-ExM images, the scale bars are post-correction. (A) Infected cells were fixed and stained for CAPRIN1 (purple), dsRNA (red), N (green), and LAMP1 (yellow). The dsRNA was chosen as a counterstain to exclude the replication sites. Cells were fixed at 11 hpi. (B) Infected cells were fixed and stained for CAPRIN1 (purple), dsRNA (red), N (green), and S (yellow). Colocalization of N and S was used to identify egress compartments. Cells were fixed at 11 hpi. (C) Infected cells were processed for U-ExM and stained for CAPRIN1 (yellow), NHS (purple), and N (green). NHS was used to identify MVB-like compartments. Cells were fixed at 6 hpi.

**Supplemental Figure 2:**
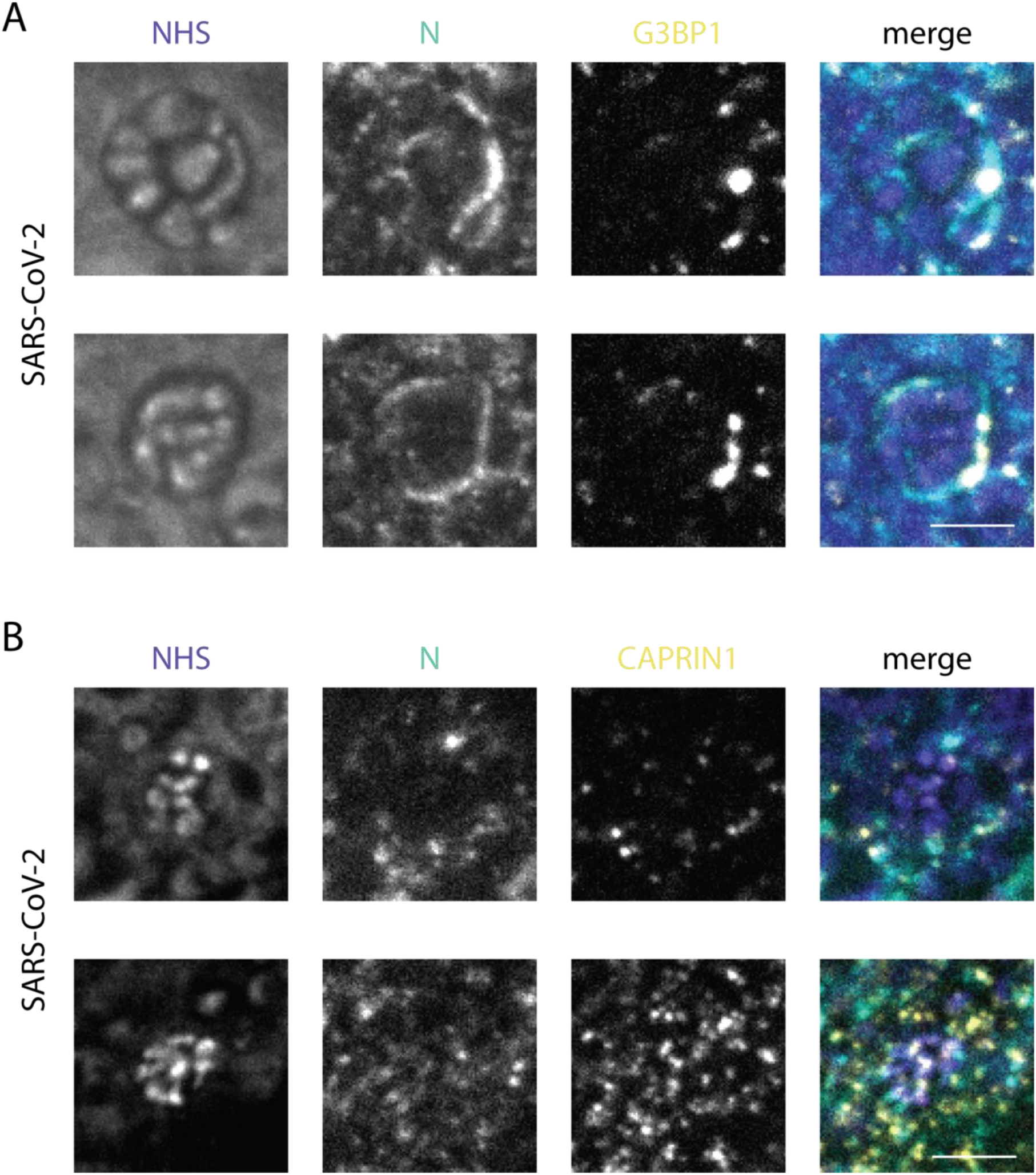
Additional U-ExM images of MVB-like compartments from SARS-CoV-2 infected cells. Additional fields of view from the samples shown in Figure 7C and Figure 8C. A549 cells were infected with SARS-CoV-2 at an MOI of 5, fixed at 6 hpi, and stained for (A) G3BP1 or (B) CAPRIN1 (yellow) and NHS (purple) and N (green). Each image is from an independent cell. Scale bars, 1 µm. For U-ExM images, the scale bars are post-correction.

**Supplemental Figure 3:**
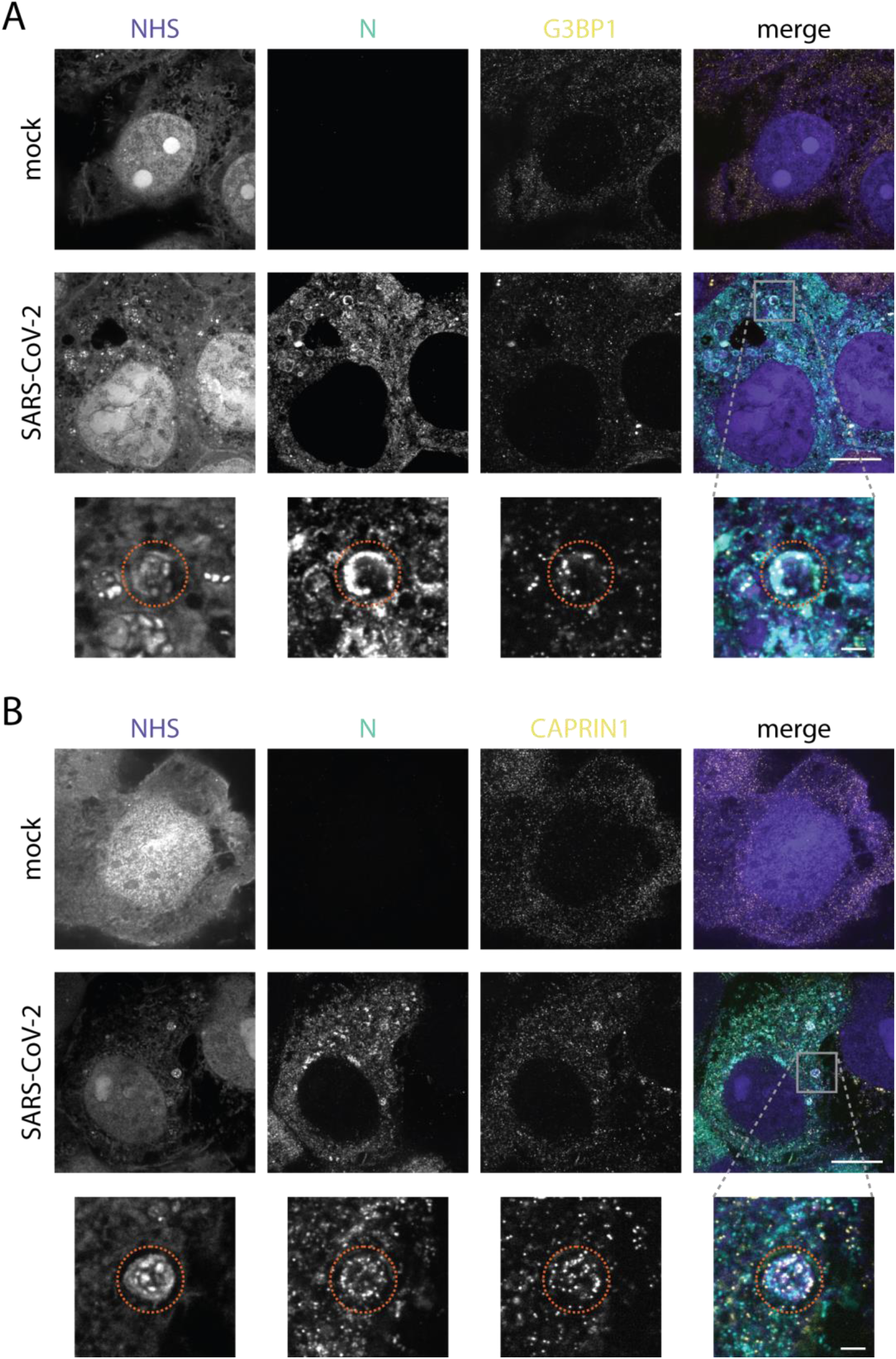
U-ExM MVB-like compartments in Vero E6 cells are comparable to those observed in A549 cells. Cells were infected with SARS-CoV-2 at an MOI of 0.1. They were fixed at 24 hpi and processed for U-ExM. Cells were stained for (A) G3BP1 or (B) CAPRIN1 (yellow), N protein (green), and NHS (purple). The MVB-like compartments are marked with a red circle. Scale bars, 10 µm; Zoom-in scale bars, 1 µm. For U-ExM images, the scale bars are post-correction.

### CAPRIN1 knockout induces aberrant cell-death morphologies and promotes syncytia formation upon SARS-CoV-2 infection

CAPRIN1 was not recruited to replication sites but was present at the MVB-like egress compartments, indicating engagement at a specific stage of the viral replication cycle. We therefore asked whether CAPRIN1 influences the course of infection and generated CAPRIN1-knockout A549 (CAPRIN1 KO) cell lines using two independent guide RNAs (gRNAs), which were validated by sequencing, Western blot, and IF (Supplemental Figure 4).

In uninfected cells, the morphology of CAPRIN1 KO cells was comparable to that of wild type (WT) cells. G3BP1 displayed a diffuse cytoplasmic distribution with scattered puncta, indicating that loss of CAPRIN1 did not promote spontaneous SG assembly (Figure 9A). This is consistent with the proposed role of CAPRIN1 as a promoter of SG formation^21,22^.

**Figure 9:**
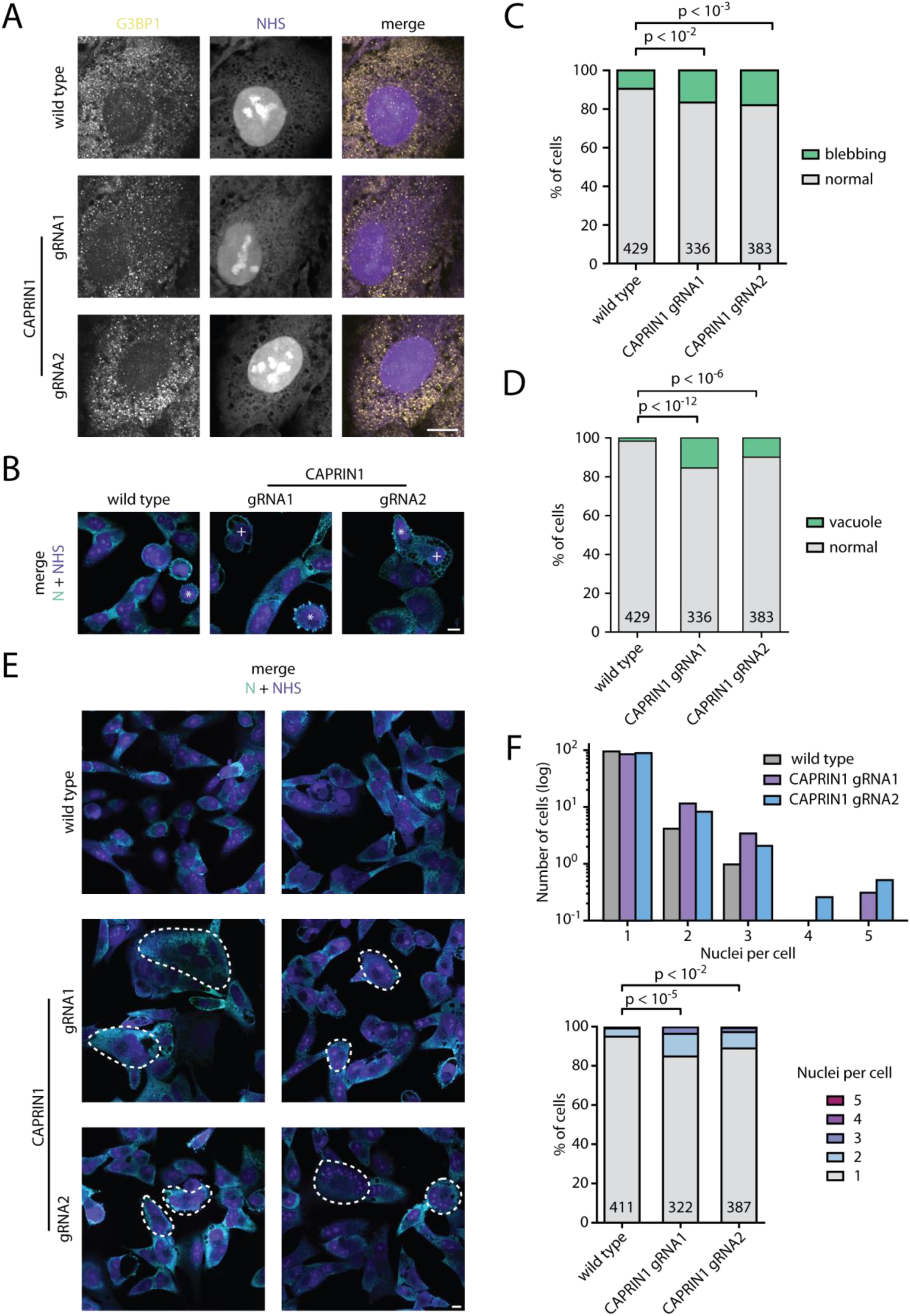
CAPRIN1 KO enhances SARS-CoV-2-induced cell death morphologies and syncytia formation. A549 cells (WT) and CAPRIN1 KO cells (gRNA1 or gRNA2) were infected with SARS-CoV-2 at an MOI of 0.1 and fixed at 24 hpi for IF. Scale bars, 10 µm. (A) Uninfected cells were stained for G3BP1 (yellow) and counterstained with NHS (purple). (B) Infected cells were stained for N (green) and NHS (purple) and scored for death morphology. Two phenotypes were counted: cells with membrane-blebbing (marked with a white *) and cells with large cytoplasmic vacuoles (marked with a white +). (C) Quantification of the fraction of cells displaying the membrane-blebbing phenotype in WT and CAPRIN1 KO cells. The p-values from a Fisher’s exact test are shown; the number of cells counted per condition is shown on the bar graph. (D) Quantification of the fraction of cells displaying large cytoplasmic vacuoles in WT and CAPRIN1 KO cells. The p-values from a Fisher’s exact test are shown; the number of cells counted per condition is shown on the bar graph. (E) Infected cells were stained for N (green) and NHS (purple) and scored for syncytia. (F) Quantification of the number of nuclei per cell in WT and CAPRIN1 KO cells. This is shown as absolute counts (upper) and fraction of total (lower). The p-values from a Mann-Whitney U test (Wilcoxon rank-sum) are shown; the number of cells counted per condition is shown on the bar graph.

Upon SARS-CoV-2 infection, both CAPRIN1 KO cell lines showed an increased cytopathic effect (CPE) (Figure 9B). In WT cells, a fraction exhibited morphological features consistent with classical apoptosis: cell shrinkage, reduction of the cytoplasmic area, and membrane blebbing^59,60^. This apoptotic phenotype was also present in the CAPRIN1 KO cells, but with higher frequency. The blebbing phenotype occurred in 17-18% of CAPRIN1 KO cells compared to 10% in WT cells (Figure 9B and C). A second prominently occurring death phenotype in CAPRIN1 KO cells was the appearance of large vacuoles occupying most of the cytoplasmic volume. These vacuoles had low NHS staining, indicating that they were protein-depleted and fluid-filled. Morphologically, they resembled the vacuoles described for methuosis, a non-apoptotic cell death pathway^61^. The vacuole phenotype appeared in 10-16% of CAPRIN1 KO cells compared to 2% of the WT cells (Figure 9B and D). Since this morphology is distinct from classical apoptosis^59^, CAPRIN1 KO cells appear to die, at least in part, through a non-apoptotic route.

In addition to the altered death morphology, CAPRIN1 KO cells displayed a higher frequency of syncytia formation compared to WT cells (Figure 9E). Syncytia were observed in 11-15% of KO cells and were less frequent in WT cells (5%). Most syncytia contained two nuclei, with the highest observed number being five nuclei per cell in both CAPRIN1 KO cell lines (Figure 9F). Together, the altered death morphology and increased syncytia formation appeared only upon infection, indicating an infection-dependent role for CAPRIN1.

**Supplemental Figure 4:**
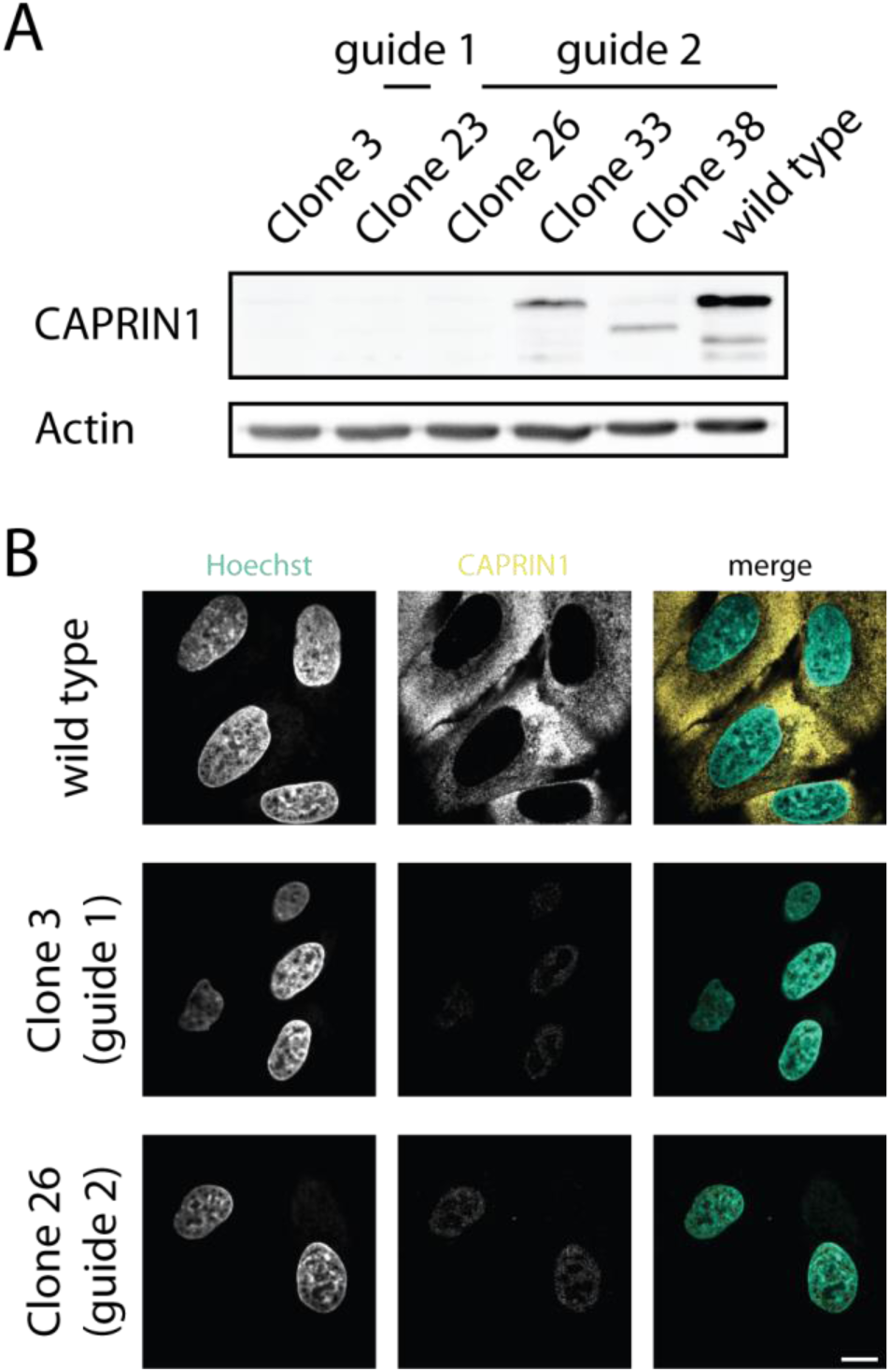
Validation of the CAPRIN1 KO cell lines by WB and IF. A549 cells were used to generate CRISPR/Cas9-mediated CAPRIN1 KO cells. (A) Western blot of five candidate clones generated from two different gRNAs. Clones 3 (gRNA1) and 26 (gRNA2) were chosen for use in this study. β-actin served as a loading control. (B) WT and CAPRIN1 KO cells were stained for CAPRIN1 (yellow) and Hoechst 33342 (green). Scale bar, 10 µm.

## Discussion

While G3BP1 is the best-characterized SG interactor of the SARS-CoV-2 N protein, the wider N protein interactome remains largely unexplored. This raises three questions: whether other stress granule proteins that are connected to G3BP1 also influence infection, where they localize in the viral cycle, and what functional consequences their engagement carries. Here we address these questions for an underexplored candidate, CAPRIN1. We show that it localizes to the viral egress compartments, which we identify as LAMP1-positive CD63-enriched MVB-like compartments, and that the loss of CAPRIN1 enhances infection-induced cell death with a distinct death morphology and syncytia formation.

Locating CAPRIN1 at egress sites prompted us to examine the identity of these compartments more closely. The current model of β-coronavirus egress proposes the use of LAMP1-positive lysosomes containing multiple virions for exocytic release^5^. We combined LAMP1 and CD63 labelling with the ultrastructural morphology from NHS staining and determined that the egress compartments are more consistent with late-endosomal/MVB-like compartments. These observations refine the current model of egress of SARS-CoV-2. Support for this reinterpretation comes from two lines of evidence: the limited specificity of the markers used in earlier work, as LAMP1, Rab7, and cathepsin D localize to both lysosomes and late endosomes/MVBs^62,63^, and the ultrastructure of egress-associated compartments, which Walia *et al.* (2024)^8^ described as multivesicular and multilamellar by TEM. Previous work^8^ demonstrated that ORF3a both hyperactivates Rab7, which leads to enhanced endosomal maturation and thus increased MVB formation, and inhibits the HOPS complex^6,7^, which blocks lysosome fusion^64,65^. This activation of an MVB-upstream process and inhibition of an MVB-downstream process would result in an increase in MVB exocytosis^66^ and likely release of infectious virus from MVB-like compartments.

Importantly, the two models of egress through lysosomes versus MVB-like compartments are not mutually exclusive (Figure 10). Walia *et al.* (2024)^8^ provide evidence for the presence of both pathways. They showed that the upstream Rab7 hyperactivation and downstream inhibition of the HOPS-complex produce two populations: 1) enlarged Rab7-positive N-containing compartments that fail to acquire Arl8b at the late-endosomal/MVB stage, which may correspond to the MVB-like compartments in step 4a of Figure 10, and 2) an Arl8b-positive and LAMP1-positive mature lysosome population proposed to undergo exocytosis, corresponding to step 4b of Figure 10. The distinction between these two compartments matters because they rely on two different release machineries. Arl8b-dependent lysosomal exocytosis proceeds through the BORC-Arl8b axis and its associated SNAREs^5,7^, whereas MVB-plasma membrane fusion follows the Rab27-dependent exosomal pathway, engaging distinct SNARE machinery^63,67^. Based on the detection of CD63, we propose that these compartments engage the MVB-associated exosomal machinery^68^, giving rise to an additional egress route alongside the lysosomal one.

**Figure 10:**
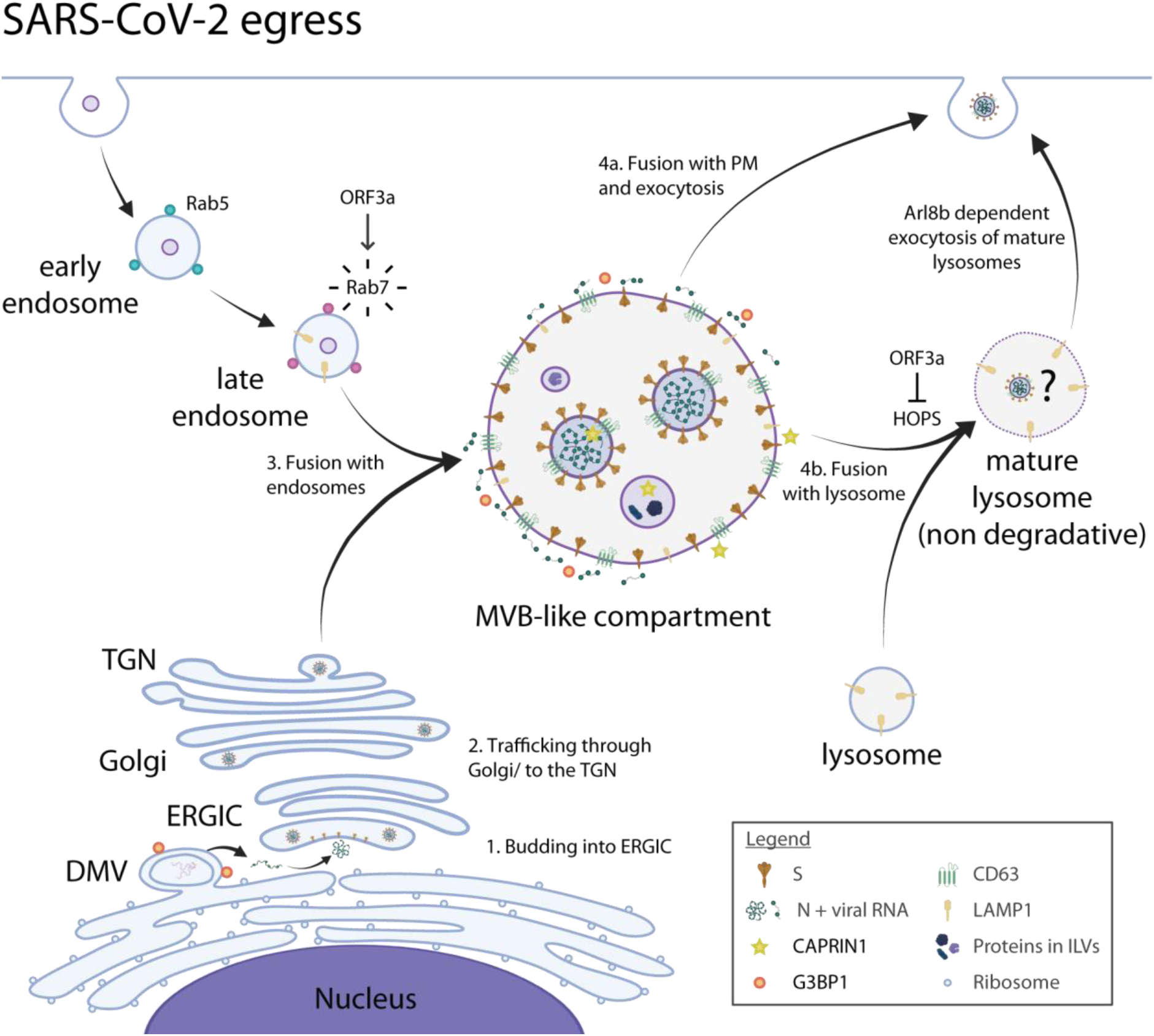
SARS-CoV-2 egress through MVB-like compartments or lysosomes. Viral genome replication occurs in DMVs that are derived from the rough ER. G3BP1 is present at the DMVs. (1) Viral RNA packaged by N protein buds into the lumen of the ERGIC, acquiring its lipid envelope with the structural proteins S, M and E. (2) Assembled virions are transported through the Golgi apparatus to the trans-Golgi network (TGN). (3) Vesicles containing virions fuse with endosomes to form MVB-like compartments. These compartments contain G3BP1 and CAPRIN1 on the limiting membrane, with CAPRIN1 also associated with the ILVs. At the same time, ORF3a activates Rab7 to drive endosome maturation. (4a) Based on the presence of CD63 on both the limiting membrane and the ILVs, we hypothesize that the virion-containing MVB-like compartments may undergo exocytosis to release infectious virus. (4b) An alternate fate for these compartments is fusion with lysosomes to form mature lysosomes. However, lysosomal fusion is mediated by the HOPS complex, which is inhibited by ORF3a. Blocking the downstream HOPS complex would therefore enhance exocytosis directly from the MVB stage^66^, as shown in 4a. SARS-CoV-2 infection has been reported to trigger Arl8b-dependent exocytosis of mature lysosomes, which are deacidified during infection^5,7^. However, it is unclear if infectious virions are trafficked to the deacidified mature lysosomes and whether they would be released into the environment.

MVB-mediated egress is an established strategy among enveloped viruses. In human cytomegalovirus (HCMV) infection, multiple virions are released simultaneously through fusion of CD63-positive multi-viral bodies with the plasma membrane in intermittent bulk pulses^69^, and hepatitis E virus is released within ILVs via an exosomal pathway^70^. Whether SARS-CoV-2 exits through bulk release of MVB content, exosome-associated release of individual ILVs, or a combination of both, and whether MVBs represent the main egress route, remain open questions for further studies.

Beyond the identification of MVB-like compartments, U-ExM revealed unexpected differences in the distribution of G3BP1 and CAPRIN1 at egress sites: CAPRIN1 was detected on both the limiting membrane and in the interior, whereas G3BP1 was predominantly localized on the limiting membrane. This spatial divergence, combined with the enhanced cytopathology observed upon CAPRIN1 KO, raises the question of whether CAPRIN1 acts like G3BP1 in its pro- and anti-viral roles.

G3BP1 does not fit a classical pro- or anti-viral classification during SARS-CoV-2 infection, and this is likely also the case for CAPRIN1. G3BP1 restricts viral RNA replication at replication sites but participates in budding and facilitates virion production^57,71^. We propose a similar duality for CAPRIN1: anti-viral when part of the SG machinery and limiting syncytia formation, but pro-viral when preventing CPE. The potential displacement of CAPRIN1 from G3BP1 by accumulating N protein (expected from their mutually exclusive binding to the G3BP1 NTF2-like domain^22,58^) would neutralize CAPRIN1’s contribution to SG-mediated innate immune signalling^17^, a likely anti-viral activity. Yet its complete loss exacerbates the cytopathic consequences of infection: CAPRIN1 KO cells die more frequently through a non-apoptotic route and form more syncytia, which is an S-driven pathology that facilitates cell-to-cell spread^72,73^ and is linked to severe COVID-19^74,75^. The formation of syncytia has not previously been linked to the loss of an SG protein, and how CAPRIN1 restrains this process remains unclear. Because CAPRIN1 accumulates at the MVB-like compartments in which S is also present, one possibility is that its loss alters S protein trafficking. It remains open whether this protective effect is specific to SARS-CoV-2 or a general function of CAPRIN1 in limiting infection-induced cell death.

This study provides the first high-spatial resolution and functional data on CAPRIN1 in the context of SARS-CoV-2 infection. By using the increased resolution from U-ExM, we could demonstrate the localization of CAPRIN1 to the SARS-CoV-2 egress compartments and further characterize the nature of viral egress. Collectively, we show the distinct behaviour of two SG proteins, which in the context of SARS-CoV-2 infection appear to have activities beyond a canonical SG function.

## Acknowledgements

This research was supported by the Maxwell computational resources operated at Deutsches Elektronen-Synchrotron DESY, Hamburg, Germany and the computing resources at Medizinische Hochschule Hannover (MHH) Information Technology (MIT) High Performance Computing (HPC) team, Hannover, Germany. We thank the Advanced Light and Fluorescence Microscopy (ALFM) Facility at the Centre for Structural Systems Biology (CSSB). Flow cytometry was performed in the FACS core facility at the Leibniz Institute of Virology (LIV). We would like to thank Arne Düsedau for his technical and practical expertise and support. The SP, CU, and JBB labs were funded by RTG 2771 Humans and Microbes project no. 453548970, and the CU and JBB labs were additionally funded by RTG 2887 VISION project number 497350882. The Bosse lab was also funded by the Deutsche Forschungsgemeinschaft (DFG, German Research Foundation) under Germany’s Excellence Strategy - EXC 2155 - project number 390874280, the CRC 1648 Emerging Viruses project number SFB 1648/1 2024–512741711, the Research Unit FOR5200 DEEP-DV (443644894) project BO 4158/5-1 and BO 4158/5-2, the Research Unit FOR5898 AdBHealth (548065690) project BO 4158/9-1, and the German Center for Infection Research (DZIF) grants IICH TTU 07.918, IICH TTU 07.861 and IICH TTU 07.863 as well as the Wellcome Trust through a Collaborative Award (209250/Z/17/Z), Behörde für Wissenschaft, Forschung und Gleichstellung (BWFG) Hamburg through “Hamburg-X Infektionsforschung”, and the Leibniz ScienceCampus InterACt, funded by the BWFG Hamburg and the Leibniz Association (W75/2022). We thank Krzysztof Pyrć, Toni Luise Meister, and Stephanie Pfänder for the A549-ACE2-TMPRSS2 cells.

## Author contributions

JSW: Conceptualization, Formal Analysis, Investigation, Methodology, Validation, Visualization, Writing – Original Draft Preparation, Writing – Review & Editing; JS: Investigation, Validation, Visualization, Writing – Review & Editing; JMB: Investigation, Resources, Validation, Writing – Review & Editing; EK: Investigation, Validation, Writing – Review & Editing; RS: Investigation, Validation, Visualization; SP: Investigation, Methodology, Funding Acquisition, Supervision, Writing – Review & Editing; CU: Funding Acquisition, Supervision, Writing – Review & Editing; TKS: Conceptualization, Formal Analysis, Investigation, Methodology, Project Administration, Resources, Supervision, Validation, Visualization, Writing – Original Draft Preparation, Writing – Review & Editing; JBB: Conceptualization, Funding Acquisition, Methodology, Project Administration, Supervision, Writing – Review & Editing

## Declaration of generative AI and AI-assisted technologies in the writing process

During the preparation of this work, the authors used Opus 5.0/Anthropic and Grammarly to improve the manuscript’s readability and language. After using these tools, the authors reviewed and edited the content as needed and take full responsibility for the published article.

## Notes

### Competing Interest Statement

The authors have declared no competing interest.

